# Single-Cell Transcriptomics Unveils Skin Cell Specific Antifungal Immune Responses and IL-1Ra-IL-1R Immune Evasion Strategies of Emerging Fungal Pathogen *Candida auris*

**DOI:** 10.1101/2024.10.22.619653

**Authors:** Abishek Balakumar, Diprasom Das, Abhishek Datta, Abtar Mishra, Garrett Bryak, Shrihari M Ganesh, Mihai G. Netea, Vinod Kumar, Michail S Lionakis, Devender Arora, Jyothi Thimmapuram, Shankar Thangamani

## Abstract

*Candida auris* is an emerging multidrug-resistant fungal pathogen that preferentially colonizes and persists in skin tissue, yet the host immune factors that regulate the skin colonization of *C. auris in vivo* are unknown. In this study, we employed unbiased single-cell transcriptomics of murine skin infected with *C. auris* to understand the cell type-specific immune response to *C. auris. C. auris* skin infection results in the accumulation of immune cells such as neutrophils, inflammatory monocytes, macrophages, dendritic cells, T cells, and NK cells at the site of infection. We identified fibroblasts as a major non-immune cell accumulated in the *C. auris* infected skin tissue. The comprehensive single-cell profiling revealed the transcriptomic signatures in cytokines, chemokines, host receptors (TLRs, C-type lectin receptors, NOD receptors), antimicrobial peptides, and immune signaling pathways in individual immune and non-immune cells during *C. auris* skin infection. Our analysis revealed that *C. auris* infection upregulates the expression of the IL-1RN gene (encoding IL-1R antagonist protein) in different cell types. We found IL-1Ra produced by macrophages during *C. auris* skin infection decreases the killing activity of neutrophils. Furthermore, *C. auris* uses a unique cell wall mannan outer layer to evade IL-1R-signaling mediated host defense. Collectively, our single-cell RNA seq profiling identified the transcriptomic signatures in immune and non-immune cells during *C. auris* skin infection. Our results demonstrate the IL-1Ra and IL-1R-mediated immune evasion mechanisms employed by *C. auris* to persist in the skin. These results enhance our understanding of host defense and immune evasion mechanisms during *C. auris* skin infection and identify potential targets for novel antifungal therapeutics.

## Introduction

*C. auris* was recently categorized as an urgent threat by the US Centers for Disease Control and Prevention (CDC) and classified in the critical priority fungal pathogens group by the World Health Organization (WHO) [1–3]. Unlike other *Candida* species, such as *Candida albicans,* which colonizes the gastrointestinal tract, *C. auris* preferentially colonizes the human skin, leading to nosocomial transmission and outbreaks of systemic fungal infections [4–6]. Furthermore, unlike skin-tropic fungal pathogens such as *Malassezia* [7], *C. auris* not only colonizes the epidermis of the skin but also enters the deeper dermis, a phenomenon that was not observed previously [4]. *C. auris* can persist in skin tissues for several months and evade routine clinical surveillance [4, 8]. Given that the majority of *C. auris* isolates exhibit resistance to several FDA-approved antifungal drugs, a deeper understanding of *C. auris*-host interactions is critical to understanding the pathogenesis and developing potential new host-directed therapeutic approaches to prevent and treat this newly emerging skin tropic fungal pathogen. Recent evidence from our laboratory indicates that *C. auris* skin infection leads to fungal dissemination, suggesting skin infection is a source of invasive fungal infection [9]. Because skin infection is a prerequisite for *C. auris* transmission and subsequent invasive disease, understanding the immune factors involved in skin defense against *C. auris* is important to understand the pathogenesis of this skin tropic fungal pathogen. Though antifungal host defense mechanisms against oral, gut, vaginal, and systemic infections of *C. albicans* are well known [10, 11], to date, almost nothing is known regarding the skin immune responses against *C. auris*.

The fungal cell wall components represent the predominant pathogen-associated molecular patterns (PAMPs) directly interacting with the host to orchestrate the antifungal immune response [12]. Recent evidence indicates that the cell wall of *C. auris* is structurally and biologically unique compared to other *Candida* species, including *C. albicans* [13]. The outer cell wall mannan layer in *C. auris* is highly enriched in β-1,2-linkages and contains two unique Mα1-phosphate side chains not found in other *Candida* species [13]. *C. auris* differentially stimulates cytokine production in peripheral blood mononuclear cells and has a more potent binding to IgG than *C. albicans* [14, 15]. Given that *C. auris* possesses a unique outer cell wall layer and preferentially colonizes and persists in skin tissue long-term, understanding the skin immune responses during the dynamic pathogen infection *in vivo* is very important but has not been explored so far. Furthermore, classical population-based gene expression studies using *in-vitro* differentiated immune cells and mouse models of systemic infection do not completely represent the host defense mechanisms against skin infection *in vivo.* In addition, understanding the skin immune responses against *C. auris* is critical, as the murine skin model is widely used to study disease pathogenesis and *C. auris*-host interactions [4, 16–20]. A closer look into cell type-specific host responses requires single-cell resolution to encompass all cell types, including immune and non-immune cells involved in host defense against *C. auris* skin infection *in vivo*.

To comprehensively define the transcriptome profiling of mouse skin infected with *C. auris in vivo* at single-cell resolution, we employed unbiased single-cell RNA sequencing (scRNA-Seq) profiling in skin tissues collected from uninfected and *C. auris*-infected mice. Our scRNA-Seq analysis identified immune cells such as neutrophils, inflammatory monocytes, macrophages, dendritic cells, T cells, NK cells, and non-immune cells such as fibroblasts accumulated at the site of infection. The scRNA-Seq revealed how skin reprograms genes and signaling pathways in immune and non-immune cell types following *C. auris* infection. The comprehensive transcriptomic profiling identified the transcriptional changes in genes that encode cytokines, chemokines, host receptors (TLRs, C-type lectin receptors, NOD receptors), antimicrobial peptides, and signaling pathways upregulated in individual myeloid cells, T cells, NK cells, fibroblast, and other non-immune cells. We identified the upregulation of the IL-1RN gene (encoding IL-1R antagonist protein) in different cell types during *C. auris* skin infection. Subsequently, using mouse models of *C. auris* skin infection and immune cell depletion studies, we elucidated the role of IL-1Ra in *C. auris* skin infection. We observed that the IL-1Ra level was significantly increased in the *C. auris*-infected skin tissue compared with *C. albicans*. *C. auris* infection induces IL-1Ra in macrophages and decreases the killing activity of neutrophils. Furthermore, *C. auris* evades IL-1R-mediated host defense through a unique outer mannan layer to persist the skin tissue.

Collectively, this study, for the first time, identified the previously unknown immune and non-immune cell type-specific skin responses and molecular events of skin-*C. auris* interactions *in vivo* at single-cell resolution. Furthermore, we demonstrated the IL-1Ra and IL-1R-mediated immune evasion mechanisms employed by *C. auris* to persist in the skin. This knowledge will be instrumental in understanding the host-pathogen interactions of *C. auris.* and will form a strong platform for developing novel host and pathogen-directed antifungal therapeutic approaches that potentially target IL-1Ra and fungal mannan, respectively.

## Results

### Unbiased scRNA seq profiling identified phagocytic cells, dendritic cells, T cells, NK cells, and fibroblast accumulated at the site of *C. auris* skin infection *in vivo*

To identify the immune and non-immune cells involved in host defense against *C. auris* skin infection, we performed unbiased scRNA seq profiling from skin tissues collected from uninfected and *C. auris*-infected mice. A group of mice was infected intradermally with 1-2 × 10^6^ CFU of *C. auris*, and another group injected with 100 µl PBS was used as an uninfected control group (Figure 1A). To capture the transcriptome profile of both innate and adaptive immune responses during *C. auris* skin infection *in vivo*, we have chosen 12 days after infection for our analysis. After 12 days post-infection (DPI), skin tissues from uninfected and infected groups (3 mice per group) were collected, minced, and digested to make single-cell suspensions as described. The single-cell suspension from infected and uninfected groups was subjected to single-cell partitioning, and RNA was sequenced using the droplet-based 10X Genomics Chromium platform (Figure 1A). After quality filtering by removing noise and batch effects, our data comprised 70,350 cells (Table S1). The ambient RNA from the data was eliminated, and the expression level of individual genes was quantified based on the number of UMIs (unique molecular indices) detected in each cell (Table S2). The alignment of the sequencing reads to the mouse reference genome resulted in the overall coverage of 32,285 genes.

**Figure 1:**
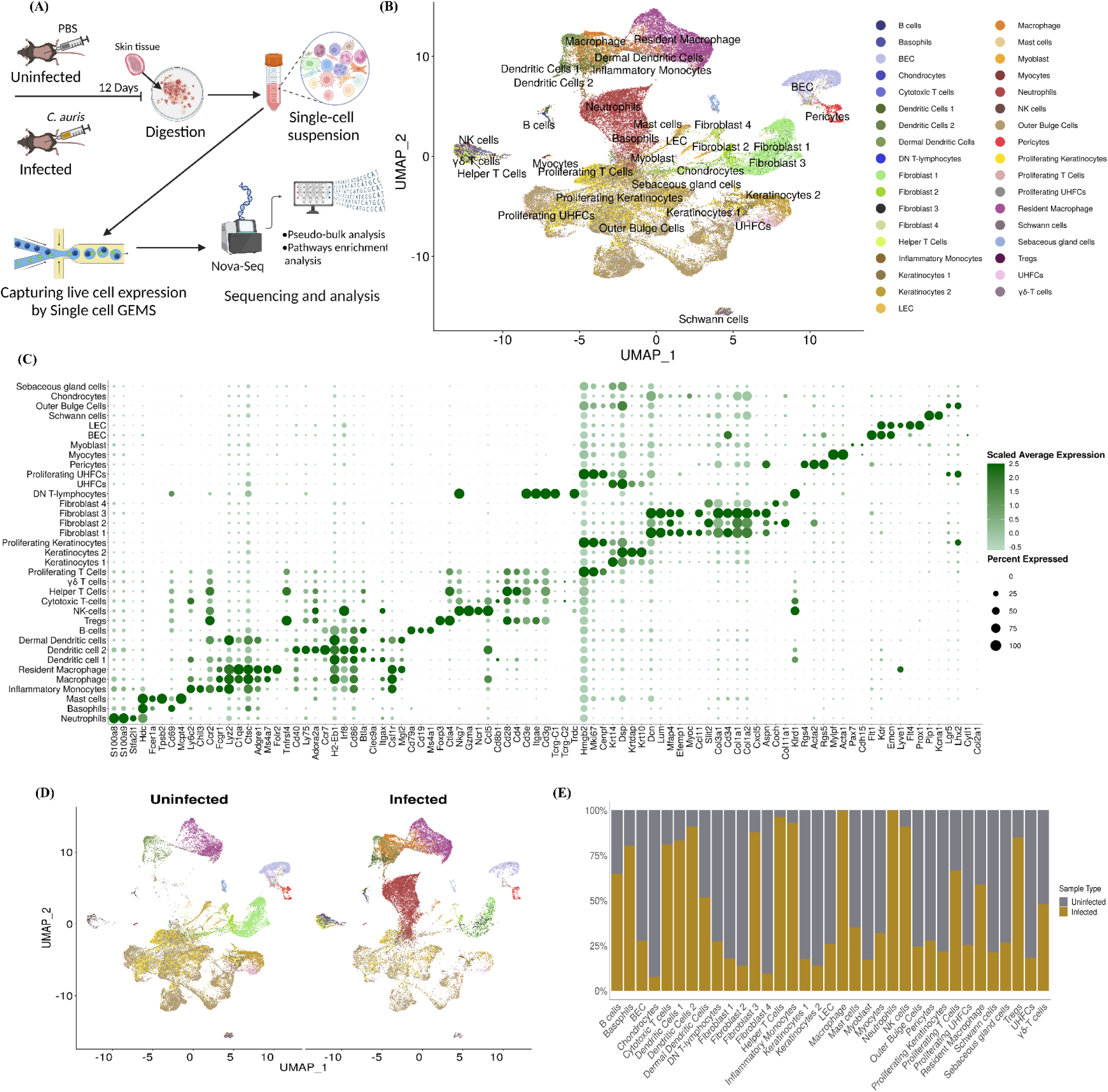
Unbiased scRNA seq profiling identified phagocytic cells, dendritic cells, T cells, NK cells, and fibroblast accumulated at the site of *C. auris* skin infection *in vivo*. A) Schematic representation of the study groups, infection course, sample processing, and single-cell preparation for scRNA sequencing. B) Uniform Manifold Approximation and Projection (UMAP) of identified cell types in the murine skin after clustering. Each cluster was assigned as an individual cell type. After identification, 35 cell types were assigned as individual clusters. C) The dot plot represents the canonical gene marker signatures to classify cell types based on specific identities. A range of 2-7 marker gene expressions were assigned to identify a cell type in the cluster. D) The UMAP of uninfected and *C. auris* infected murine skin tissue with resident and recruited cell population depicting the cell type heterogenicity between the two groups. E) The composition of the individual cell types in the uninfected and *C. auris* infected murine skin. Cell type proportions were normalized from the total cells detected in each sample.

After unsupervised graph-based clustering and reference-based annotation from the dataset GSE181720, 30 clusters were identified in our scRNA seq dataset, and cell-type identity was assigned to all the cell populations in the clusters (Figure 1B). To identify each cell lineage from 35 distinct cell types in our scRNA seq dataset, cell-type specific identity was assigned based on the canonical marker expression described elsewhere [21–23] (Figure 1C). After cell-type annotation, we identified 16 cell types as immune cells and the rest 19 as non-immune cells. The UMAP of *C. auris* infected samples indicated the recruitment of major immune cell types and resident cell populations (Figure 1D). Our analysis identified the percentage of immune and non-immune cells accumulated at the site of infection; (neutrophils – 99.9%, basophils – 80.5%, inflammatory monocytes – 93%, macrophages – 99.6%, dendritic cell 1 – 83.3%, dendritic cell 2 – 90.9%, helper T cell – 96.4%, cytotoxic T cells - 81%, Tregs - 84.9%, NK cells – 90.9%, B-cells - 64.7%, proliferating T cells – 66.5%, resident macrophages – 58.9%, dermal dendritic cells – 51.8%, and γδ T cells - 47.8%). Surprisingly, we identified fibroblast 3 cell types (87.9%) as major non-immune cells that showed increased accumulation in the skin tissue of infected groups (Figure 1D and 1E) (Table S1). The recruitment of neutrophils, inflammatory monocytes, macrophages, dendritic cells, and T cell subsets following *C. auris* skin infection was validated by flow cytometry (Figure S1). The proportion of cell types between the *C. auris* infected and uninfected groups varies drastically (Figure 1E) and displays cellular heterogeneity of the resident and recruited cell population. Collectively, our scRNA seq identified the accumulation of various innate and adaptive immune cells at the site of infection. In addition to phagocytic cells, DCs, and T cells, which are known to play a critical role in antifungal defense, our analysis identified an accumulation of NK cells and fibroblast during *C. auris* skin infection.

### scRNA seq revealed the host immune transcriptomic signatures in individual myeloid cells during *C. auris* skin infection

To identify the host immune genes regulated in individual myeloid cells identified during *C. auris* skin infection *in vivo*, we performed pseudo-bulk analysis by normalizing UMI counts for the target genes and identifying differentially expressed genes (DEGs) between the infected and uninfected groups. For the downstream analysis, we filtered DEGs with FDR < 5%, and the log2 fold change ≥ 2 was considered upregulated, and ≤ -2 was considered downregulated. In neutrophils, 1107 genes (1105 upregulated and 2 downregulated), inflammatory monocytes, 299 genes (217 upregulated and 82 downregulated), macrophages, 1340 genes (1322 upregulated and 18 downregulated), resident macrophages, 357 genes (143 upregulated and 214 downregulated), dendritic cell 1, 454 genes (260 upregulated and 194 downregulated), dendritic cell 2, 265 genes (244 upregulated and 21 downregulated) and dermal dendritic cells 262 (91 upregulated and 171 downregulates) were differentially expressed in the *C. auris* infected groups. The top 10 upregulated and downregulated genes in the myeloid subsets were highlighted in the volcano plot (Figure 2A). Among the top 10 DEGs in myeloid cells, chemokine genes such as *Ccl4* and *Cxcl3* in neutrophils, *Cxcl9* and *Ccl5* in inflammatory monocytes, and *Cxcr2* in dermal dendritic cells were significantly upregulated in the *C. auris* infected group (Figure 2A). In macrophages, resident macrophages, and dermal dendritic cells, *Nos2* and *Arg1* involved in nitric oxide metabolism were upregulated genes after infection. *Arg1* was also significantly upregulated in dermal dendritic cell 1 (Figure 2A). In addition, *Inhba, slpi,* and *aw112020* genes known to shift the nitric oxide metabolism were upregulated in macrophages (Figure 2A). Furthermore, the gene encoding for serum amyloid A3 protein (*Saa3)* was significantly upregulated in all the phagocytic cells (Figure S2A) (Table S10). We examined the genes that were upregulated in all phagocytic cells. *Spp1, Egln3, Slpi, Upp1, Inhba, F10, Acod1, AA467197, Ppp1r3b, Nos2, Slc2a1, Tarm1, Cd300lf,* and *Ly6a* were significantly upregulated in all the phagocytic cells during *C. auris* infection (Figure S2A) (Table S10). We identified *AA467197, Plac8, Arg1, Ccl17, Slpi,* and *AW112010* were significantly upregulated in dendritic cell 1, dendritic cell 2, and dermal dendritic cells (Figure S2B) (Table S10).

**Figure 2:**
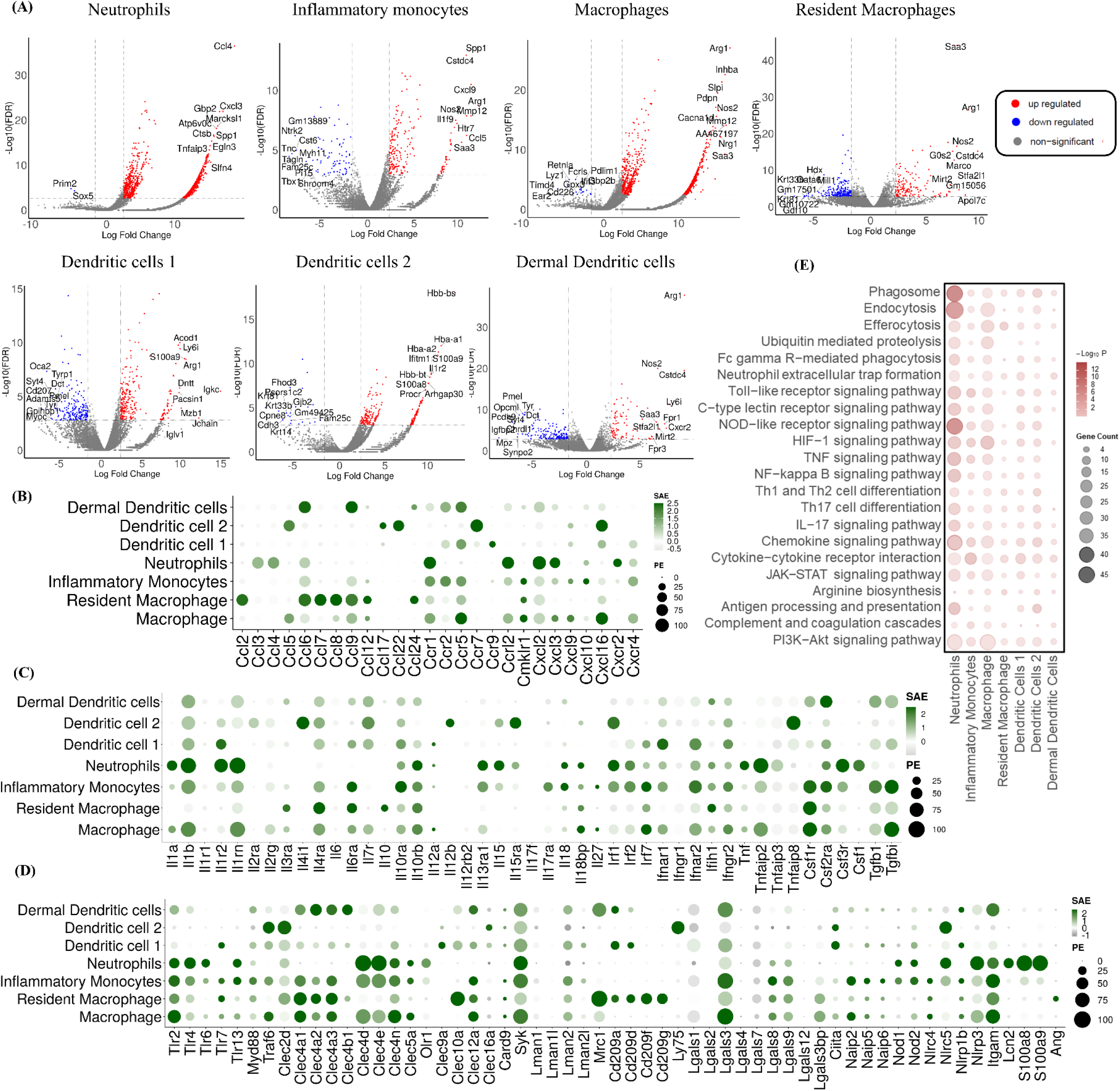
scRNA seq revealed the host immune transcriptomic signatures in individual myeloid cells during *C. auris* skin infection. A) Volcano plots indicating significant differentially expressed genes (DEGs) in the infected and uninfected neutrophils, monocytes, macrophages, resident macrophages, dendritic cell 1, dendritic cell 2 and dermal dendritic cells highlighting the top 10 upregulated and downregulated genes; The Dot plot represents the expression of selected B) chemokines and its receptors, C) cytokines and its receptors, D) fungal recognizing receptors and antimicrobial peptides (AMPs) in the myeloid subsets (Y-axis). The dot size represents the percentage of cells with expressions, and the color indicates the scaled average expression calculated from the 3 uninfected and 3 infected samples. E) The bubble plot represents the KEGG pathway of the enriched upregulated genes of the myeloid subsets (X-axis). DEGs from pseudo-bulk analysis with threshold Log 2-fold change ± 2 and FDR > 5% were considered as significant DEGs, and Log 2-fold change ≥ 2 and FDR > 5% as upregulated genes.

Next, we compared the expression of chemokines, cytokines, pattern-recognizing receptors (PRRs), and antimicrobial peptides (AMPs) among different myeloid subsets [11, 24]. The gene list from the mouse genome database was used to explore the expression of the host immune genes [25]. Among the chemokines and chemokine receptors, *Ccr1, Ccrl2, Cxcl2, Cxcl3,* and *Cxcr2* were highly expressed in neutrophils. *Ccr1, Ccr2, Ccr5,* and *Cxcr4* were highly expressed in inflammatory monocytes, whereas *Ccr1*, *Ccrl2*, *Ccr5*, *Ccl5*, *Ccl9*, *Cmklr1, Cxcl16* and *Cxcr4* were highly expressed in the inflammatory macrophage and *Ccl2, Ccl6, Ccl7, Ccl8, Ccl9, Ccl12,* and *Ccl24* were highly expressed in the resident macrophages. Dendritic cell 1 showed increased expression of *Ccr5, Ccr9,* and *Ccl16,* whereas *Ccl5*, *ccl17, Ccl22*, *Ccr7*, and *Cxcl16* were highly expressed in dendritic cell 2. *Ccl6, Ccl9, Ccr2,* and *Ccr5* were highly expressed in the dermal dendritic cells (Figure 2B). Among the cytokines and cytokine receptors, *Il1a, Il1b, Il1r2*, and *Il1rn* were highly expressed in neutrophils. *Il1a, Il1b,* and *Il1rn* were highly expressed in macrophages, whereas *Il1b* and *Il1rn* were highly expressed in monocytes and dendritic cells 1. We identified increased expression of *Il10* in the resident macrophages, *Tnf* and *Csf1* were increased in neutrophils, and *Tgfbi* expression was upregulated in monocytes and macrophages (Figure 2C).

Next, we compared the expression of TLRs, C-type lectin receptors, NOD-like receptors (NLRs), and complement receptor 3 among myeloid subsets. Neutrophils showed an increased *Tlr2, Tlr4, Tlr6,* and *Tlr13* expressions. *Tlr2, Tlr4*, *Tlr7, Tlr13,* and the adaptor molecules such as *MyD88* and *Traf6* were highly expressed in the inflammatory monocytes. *Tlr2* was highly expressed in macrophages, whereas *Tlr2* and *Tlr7* were highly expressed in resident macrophages. *Tlr7* and *Tlr2* were highly expressed in dendritic cell 1 and dermal dendritic cells, respectively. Among the C-type lectin receptors, *Clec4d* (Dectin-2), *Clec4n* (Dectin-3), and *Clec4e* (Mincle) were highly expressed in neutrophils, monocytes, and macrophages. The *Syk,* an adopter protein involved in Dectin-1 signaling, is highly expressed in all myeloid cells. *Card9*, the other downstream signaling protein of Dectin-1, is selectively expressed in all the myeloid subsets except neutrophils. The *Clec4a1, Clec4a2,* and *Clec4a3* were highly expressed in monocytes, macrophages, resident macrophages, and dermal dendritic cells, whereas the *Clec4b1* was only expressed in the dermal dendritic cells. The mannose-binding receptor *Mrc1* (Mannose receptor) was highly expressed in resident macrophages and dermal dendritic cells. The *cd209a* (DC-SIGN1), *cd209d* (SIGN-R3), *cd209f* (SIGN-R8), and *cd209g* encoding DC-SIGN were highly expressed in resident macrophages. The *Lgals3* (Galectin-3) was highly expressed in monocytes and macrophages, followed by dermal dendritic cells and dendritic cell 1. We identified increased expression of intracellular PRR, *Nod1,* and *Nod2* in the neutrophils and monocytes. The *Itgam* (complement receptor 3) was highly expressed in neutrophils, monocytes, macrophages, resident macrophages, and dermal dendritic cells. We identified increased expression of *Nlrp3* in neutrophils. Among the AMPs, the *Lcn2, S100a8*, and *S100a9* were highly expressed in neutrophils, and the *Ang* was selectively expressed in the resident macrophages (Figure 2D).

To explore the enrichments of DEGs in the myeloid cells, KEGG pathway analysis was performed for significantly upregulated genes (5% FDR, Log2Fc ≥ 2) in neutrophil, monocytes, macrophages, resident macrophages, dendritic cell 1, dendritic cell 2 and dermal dendritic cells 2 to identify the upregulated pathways (Figure 2E). Among myeloid subsets, we identified several pathways involved in pathogen recognition and immune signaling that were highly enriched in neutrophils. The major innate pathways enriched in the neutrophils during *C. auris* skin infection: 1) phagocytosis (phagosome, endocytosis, efferocytosis, ubiquitin-mediated proteolysis, Fc gamma R-mediated phagocytosis, and neutrophil extracellular trap formation), 2) PRRs (Toll-like receptor signaling pathway, C-type lectin receptor signaling pathway, and NOD-like receptor signaling pathway), 3) inflammasome activation (HIF-1 signaling pathway, TNF signaling pathway, and NF-kappa B signaling pathway), 4) cytokine and chemokine signaling (chemokine signaling, cytokine-cytokine receptor pathway, and JAK-STAT signaling pathway), 5) T-helper cell (Th) differentiation (Th1 and Th2 cell differentiation, Th17 cell differentiation and IL17 signaling pathway). In addition, we identified the enriched upregulated genes in these KEGG pathways (Table S3). Collectively, our single-cell transcriptomics identified the AMPs, cytokines, chemokines, and fungal recognition receptors upregulated in myeloid subsets during *C. auris* skin infection. We identified the genes involved in nitric oxide metabolism and acute phase proteins, which were highly upregulated in different myeloid cells. Furthermore, our analysis revealed the enrichment of several pathways involved in fungal recognition, phagocytosis, cytokine and chemokine signaling, and T-cell differentiation in individual myeloid cells during *C. auris* skin infection *in vivo*.

### *C. auris* skin infection induces IL-17 and IFNγ signaling pathways in lymphoid cells

To identify the host immune genes and transcription factors (TFs) regulated in T cell subsets and NK cells during *C. auris* skin infection, we sub-clustered the lymphoid cells at 0.2 resolution to identify the subsets of CD4+ Th cells, CD8+, γδ T cells, Tregs, and NK cells (Figure 3A). Sub-clustering the lymphoid population identified 203 genes in CD4^+^ Th cells (118 upregulated and 85 downregulated), 41 genes in Tregs (34 upregulated and 7 downregulated), 18 genes in CD8^+^ cells (14 upregulated and 4 downregulated), 92 genes in γδ T cells (73 upregulated and 19 downregulated) and 16 genes in NK cells (9 upregulated and 7 downregulated) were differentially expressed, and the heatmap shows the significantly upregulated genes (log2 fold change ≥ + 2, FDR < 5% or 1%) in CD4^+^ Th cells, γδ T cells, Tregs, CD8^+^ cells, and NK cells (Figure 3D, Figure S3B and Figure S3C). We examined the genes upregulated in lymphoid subsets (Figure S3A). The *Nfkbia* was upregulated in CD4+ Th cells and CD8+ cells. *Epsti1* and *Enox2* were significantly upregulated in CD4+ Th cells and NK cells (Figure S3A) (Table S10).

**Figure 3:**
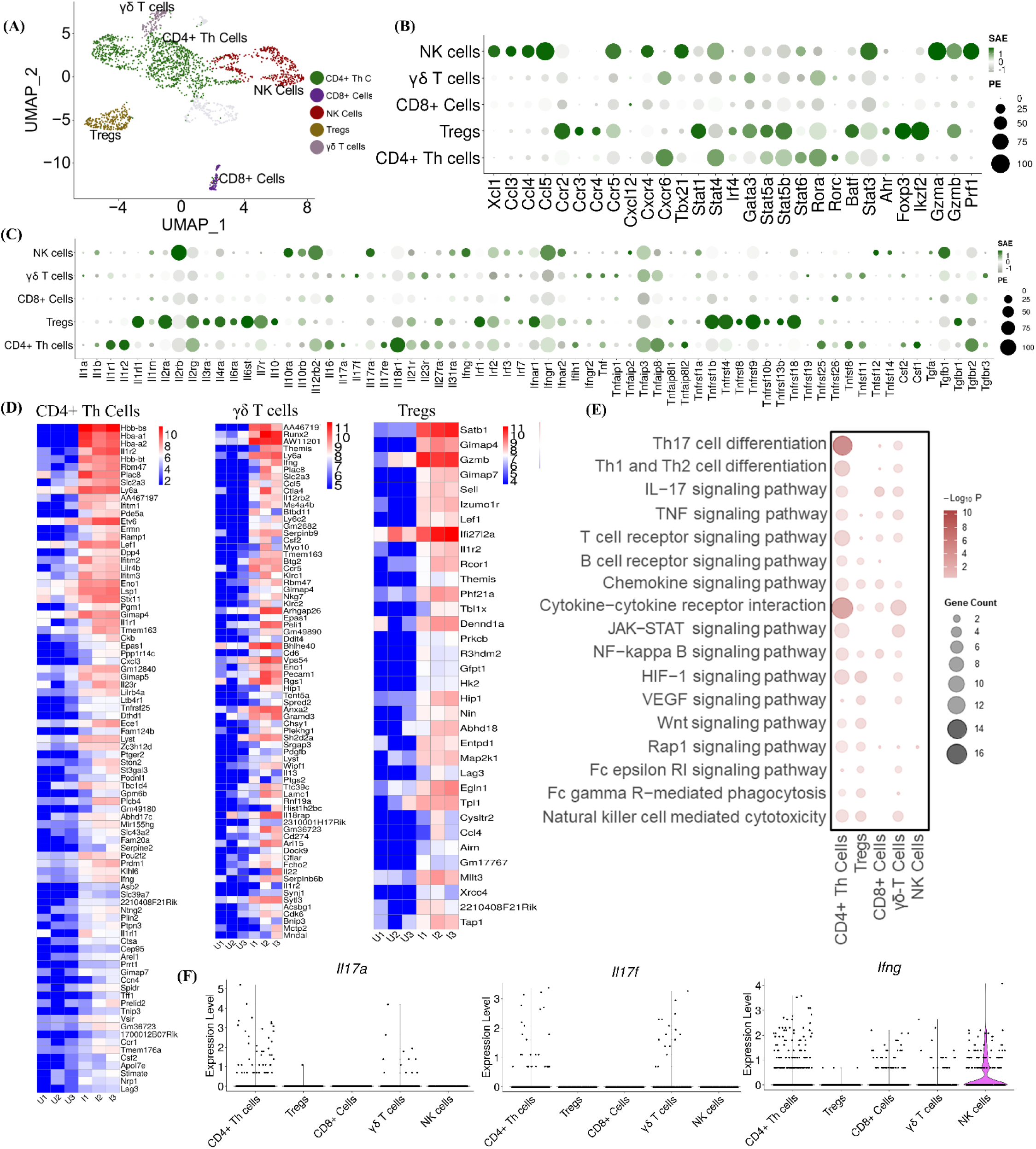
Single-cell profiling of lymphoid subsets. A) Uniform Manifold Approximation and Projection (UMAP) represents the sub-clustering analysis of lymphoid subsets at 0.2 resolution. The CD4+ Th cells, Tregs, CD8+ cells, γδ T cells, and NK cells were identified in the subpopulation. The Dot plot represents the expression of selected B) chemokines and their receptors, transcription factors (TFs), and cytolytic enzymes, and C) cytokines and their receptors in the lymphoid subsets (Y-axis). The dot size represents the percentage of cells with expressions, and the color indicates the scaled average expression calculated from the 3 uninfected and 3 infected samples. D) The heatmap represents the expression of the significantly upregulated genes in CD4+ Th cells, γδ T cells, and Tregs in uninfected and infected groups. The normalized gene counts were plotted in the heatmap, and the scale indicates red for high, blue for low, and white for moderate expression in the samples. Each column represents a different sample. E) The bubble plot represents the KEGG pathway enrichment analysis of significantly upregulated genes in the lymphoid subsets (X-axis). F) Violin plots depict the scaled count of *Il17a, Il17f*, and *Ifng* in CD4+ Th cells, Tregs, CD8+ cells, γδ T cells, and NK cells in both uninfected and infected samples. The scaled count were normalized using SCTransform method. DEGs from pseudo-bulk analysis with threshold Log 2-fold change + 1 and FDR > 5% were considered as significant upregulated genes. Upregulated genes with Log 2-fold change ≥ 2 and FDR > 1% for CD4+ Th cells and FDR > 5% for Tregs, and γδ T cells were represented for the heatmap.

Next, we compared the expression of chemokines, cytokines, and TFs among lymphoid subsets. *Cxcr6* was highly expressed in CD4^+^ Th cells and γδ T cells. We identified increased expression of *Xcl1, Ccl3, Ccl4, Ccl5, Ccr5,* and *Cxcr4* in NK cells, and *Ccr2, Ccr3,* and *Ccr4* in the Tregs. TFs such as *Stat4, Stat5a, Stat5b, Stat6, Rora, Rorc,* and *Ahr* were highly expressed in the CD4+ Th cells. Tregs showed increased expression of *Stat1, Gata3, Stat5a, Stat5b, Batf, Foxp3, Ikzf2*, *Irf4* and *Ahr*. We identified the increased *Tbx21, Stat4, Stat6,* and *Stat3* expression in NK cells. *Gzma, Gzmb,* and *prf1* in NK cells and *Gzmb* were highly expressed in the Tregs (Figure 3B). *Il16, Csf1, Il1b, Il17a, Tnf,* and *Csf2* were highly expressed in the CD4+ Th cells. Tregs showed a higher expression of *Il10*. *Il17a* and *Il17f* were selectively expressed in γδ T cells. We identified increased expression of *tgfb1* and *ifng* in the NK cells (Figure 3C).

The upregulated pathways enriched in CD4+ Th cells, CD8+ cells, γδ T cells, Tregs, and NK cells were identified from KEGG pathway analysis of the upregulated genes with Log2Fc ≥ 1 and 5% FDR threshold (Figure 3E). We identified pathways involved in Th17 cell differentiation, cytokine-cytokine receptor interaction, Th1, and Th2 cell differentiation, and HIF-1 signaling pathway were highly enriched in CD4^+^ Th cells during *C. auris* skin infection (Figure 3E). Other pathways involved in TNF, JAK-STAT, NF-kappa B and HIF-1 signaling were enriched in CD4^+^ Th cells, Tregs and γδ T cells (Figure 3E). In addition, we examined the upregulated DEGs enriched in these KEGG pathways (Table S4). Next, we used violin plots to identify the cell type level expression of *Il17a, Il17f,* and *Ifng* in CD4+ Th cells, CD8+ cells, γδ T cells, Tregs, and NK cells (Figure 3F). Our analysis revealed that *Il17a* and *Il17f* were mainly expressed in CD4^+^ Th cells and γδ T cells during *C. auris* skin infection. Our analysis identified that NK cells showed an increased expression of *Ifng*, followed by CD4^+^ Th cells, CD8^+^ cells, and γδ T cells (Figure 3F). IL-17 signaling pathway, which is known to play a critical role in antifungal defense, is upregulated in CD4^+^ Th cells and γδ T cells. Taken together, our scRNA analysis identified the expression of cytokines, chemokines, and TFs upregulated in T cells and NK cells.

### scRNA seq revealed the transcriptomic signatures in fibroblast and other non-immune cells during *C. auris* skin infection

Our unbiased scRNA profiling identified fibroblast as a major non-immune cell accumulated at the site of skin infection. To understand the transcriptomic signatures of fibroblast during *C. auris* skin infection, the gene expression profiling of fibroblast sub-clusters, fibroblast 1, fibroblast 2, fibroblast 3, and fibroblast 4, were explored. The DEGs from the pseudo-bulk analysis with a threshold of FDR < 5% and the log2 fold change ≥ 2 were used for further analysis. Log2 fold change of ≤ -2 were considered downregulated for the volcano plots. Among the DEGs in fibroblast subsets, 370 genes in fibroblast 1 (150 upregulated and 220 downregulated), 68 genes in fibroblast 2 (12 upregulated and 56 downregulated), 389 genes in fibroblast 3 (294 upregulated and 95 downregulated) and 9 genes in fibroblast 4 (1 upregulated and 8 downregulated) were regulated upon *C. auris* infection. The top 10 upregulated and downregulated genes in fibroblast subsets were denoted in the volcano plots (Figure. 4A). The antimicrobial peptide *S100a9* is the top upregulated genes in fibroblast 1, fibroblast 2 and fibroblast 4. The *S100a8*, which forms a complex with S100a9 as calprotectin, is among the top 10 upregulated genes in fibroblast 2. The serum amyloid protein *Saa3* is the highest upregulated gene in fibroblast 3, and the *Saa1* and *Saa3* were highly upregulated in fibroblast 1 (Figure. 4A). Further, the antimicrobial peptides *NOS2* and *Lcn2* are highly upregulated in fibroblast 1 and fibroblast 3 (Figure. 4A) (Figure S5A).

**Figure 4:**
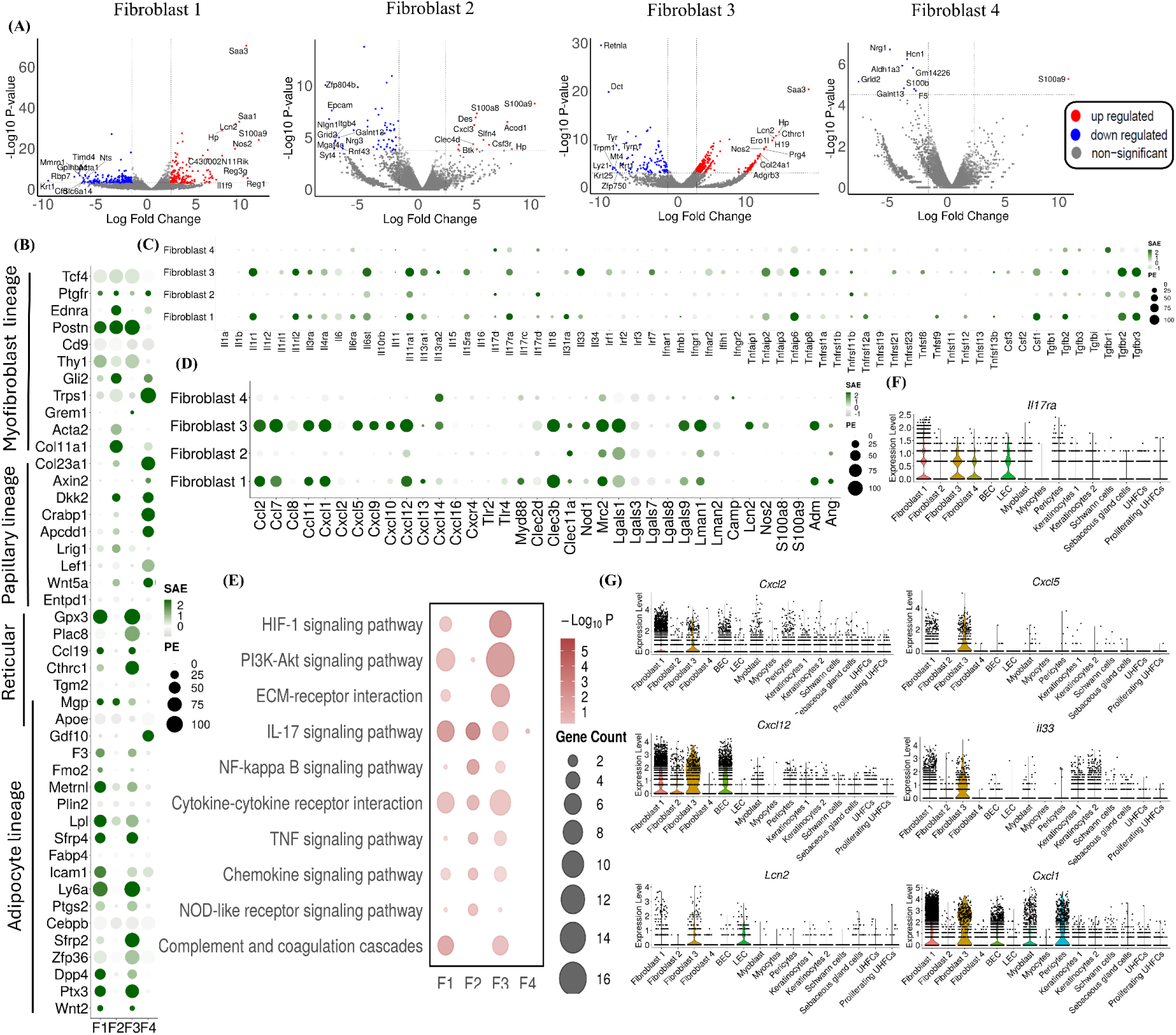
scRNA seq revealed the transcriptomic signatures in fibroblast and other non-immune cells during *C. auris* skin infection. A) The Volcano plots indicate significant differentially expressed genes (DEGs) in the infected and uninfected fibroblast subsets, highlighting the top 10 upregulated and downregulated genes. The Dot plot represents the expression of selected B) fibroblast lineage markers, C) cytokines and their receptors, and D) chemokines and their receptors, fungal recognizing receptors, and antimicrobial peptides (AMPs) in the fibroblast subsets (Y-axis). The dot size represents the percentage of cells with expressions, and the color indicates the scaled average expression calculated from the 3 uninfected and 3 infected samples. E) The bubble plot represents the KEGG pathway enrichment of the upregulated genes of fibroblast subsets (X-axis). ; Violin plots depict the scaled count of F) *IL17ra* and G) *Cxcl1, Cxcl2, Cxcl5, Cxcl12, Lcn2,* and *IL33* in fibroblast subsets and other non-immune cells in both uninfected and infected samples. The scaled count were normalized using SCTransform method; DEGs from pseudo-bulk analysis with threshold Log 2-fold change ± 2 and FDR > 5% were considered as significant DEGs and Log 2-fold change ≥ 2 and FDR > 5% as upregulated genes.

We classified the fibroblast subsets based on their lineage-specific marker expression. Fibroblasts 1 and 3 highly expressed most adipocyte lineage markers, and fibroblast 4 substantially expressed papillary lineage markers (Figure 4B). Next, we compared the expression of chemokines, cytokines, fungal-recognizing receptors, and AMP among four fibroblast subsets (Figures 4C and 4D). *Il33, Csf1*, and *Tgfb2* were highly expressed in fibroblast 3 and fibroblast 1, whereas *Il17d* was selectively expressed in fibroblast 2 and fibroblast 4. Among the cytokine receptors, *Il1r1, Il1rl2, Il6st, Il11ra1, Il17ra, Tgfbr2,* and *Tgfbr3* were highly expressed in fibroblast 3 and fibroblast 1, whereas *Il11ra1* and *Tgfbr3* were highly expressed in fibroblast 2 (Figure 4C). Among the chemokines, *Ccl2, Ccl7, Ccl11, Cxcl1, and Cxcl12* were highly expressed in fibroblast 1 and fibroblast 3. *Cxcl5, Cxcl9, Cxcl10,* and *Cxcl14* were highly expressed in fibroblast 3. Among the fungal recognizing receptors, *Clec3b, Mrc2, Lgals1, Lgals9, Lman1,* and *Clec11a* were highly in fibroblast 3 and fibroblast 1 (Figure 4D). We identified increased expressions of *Clec11a, Mrc2, Lgals1,* and *Lman1* in fibroblast 2 and selective expression of *Nod2* in fibroblast 3. Among the AMPs, fibroblast 3 showed increased expression of *Lcn2, Nos2,* and *Amd,* whereas *Amd* and *Ang* were highly expressed in fibroblast 1 (Figure 4D).

To identify the upregulated pathways in fibroblast subsets during *C. auris* skin infection, KEGG pathway enrichment was performed for upregulated genes with Log2Fc ≥ 2 and 5% FDR in fibroblast 1, fibroblast 2, fibroblast 3, and fibroblast 4 (Figure 4E). We identified pathways involved in HIF-1 signaling, PI3K-Akt signaling, cytokine-chemokine receptor interaction, and complement coagulation cascade pathways that were highly enriched in fibroblast 3 and fibroblast 1. ECM-receptor interaction pathway was enriched in fibroblast 3. The IL-17 signaling pathway was enriched in all three fibroblast subtypes except fibroblast 4 (Figure 4E). Next, we analyzed other non-immune cells, such as BEC, outer bulge cells, and pericytes, that showed DEGs during *C. auris* skin infection (Figure S4). We examined the upregulated DEGs enriched in these KEGG pathways (Table S5). KEGG pathways, such as Th17 cell differentiation and cytokine-cytokine receptor interactions, were highly enriched in BEC, pericytes, and outer bulge cells (Figure S5B-5D). To understand if fibroblast and other non-immune skin cells play a role in the recruitment of immune cells during *C. auris* infection, we analyzed the expression of genes such as *IL-17ra* and chemokines involved in recruiting immune cells such as neutrophils. Among non-immune cells, fibroblast subsets and LEC showed increased expression of the *IL-17ra* gene (Figure 4E). The chemoattractants such as *Cxcl1, Cxcl2, Cxcl5, Cxcl12, Lcn2*, and *Il33* involved in the recruitment of neutrophils were highly expressed in different non-immune cells, but fibroblast subsets showed relatively higher expression (Figure 4G). Taken together, our unbiased scRNA seq revealed the increased expression of cytokines, chemokines, PRRs, and AMPs in fibroblasts and other non-immune cells that could either directly (or) indirectly contribute to the host defense against *C. auris* skin infection.

### *C. auris* induces IL-1Ra in macrophages to decrease the neutrophil function in the skin

Our scRNA seq identified that *C. auris* skin infection upregulated the expression of the *Il1rn* gene in different cell types, especially in myeloid and lymphoid cells (Figure 5A). The *Il1rn* gene encodes four isoforms of IL-1R antagonist (IL-1Ra), which regulates the inflammatory signaling of the IL-1 pathway by competing with IL-1α and IL-1β in IL-1R activation [26, 27]. We analyzed the expression of *Il1rn* in the cell types identified in our analysis. In our dataset, the *Il1rn* was expressed only in hematopoietic cells in the *C. auris* infected samples (Figure 5A) (Figure S6A). However, in the non-hematopoietic cells, such as fibroblasts and keratinocytes, that are known to express intracellular isoforms of IL-1Ra [28, 29], the *C. auris* infection did not considerably altered *Il1rn* expression (Figure S6A).

**Figure 5.**
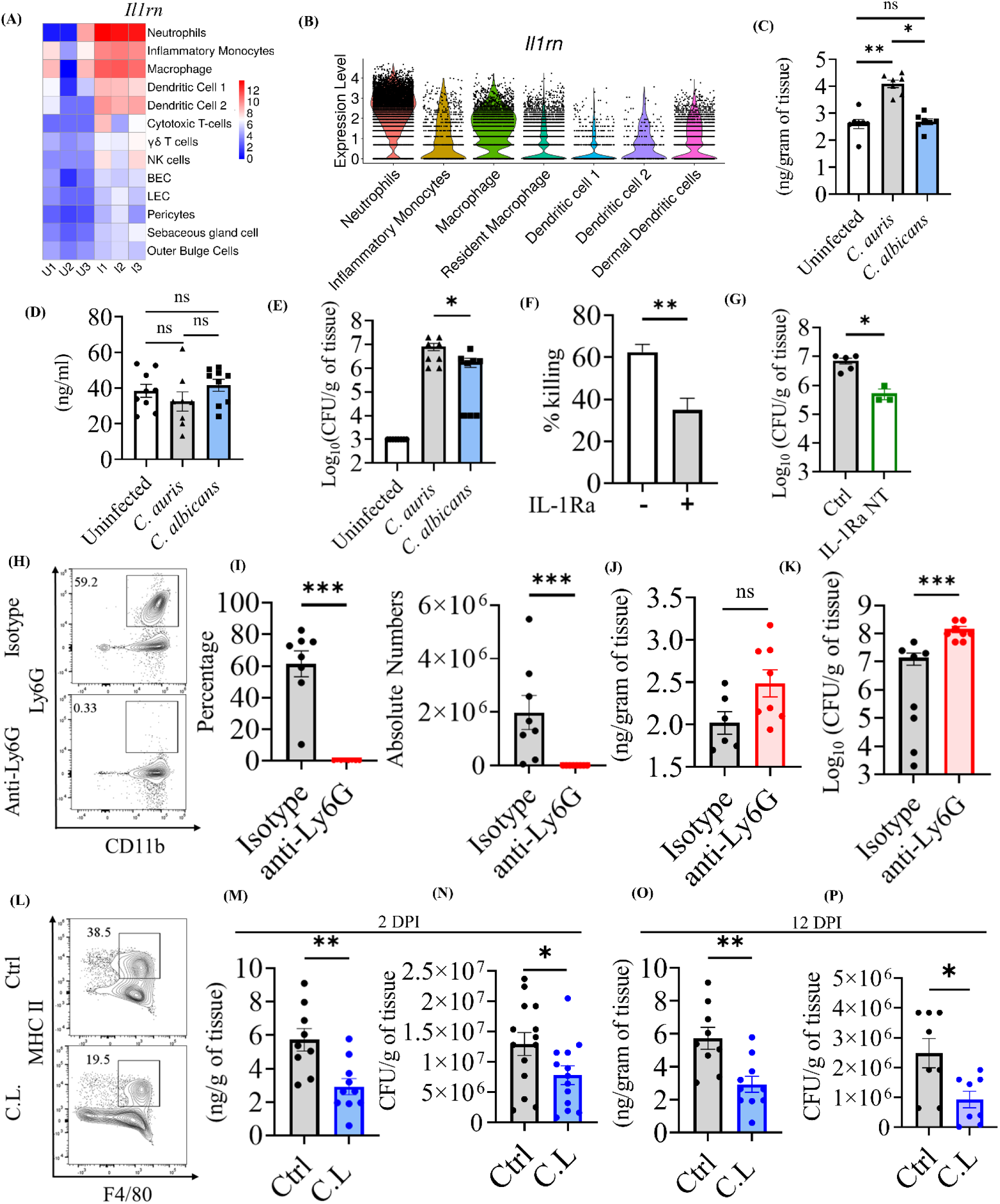
*C. auris* induces IL-1Ra in macrophages to decrease the neutrophil function in the skin. (A) Heatmap represents the expression of *Il1rn* in the cell types (y-axis) between the infected and uninfected groups. The normalized gene counts were plotted in the heatmap, and the scale indicates red for high, blue for low, and white for moderate expression in the samples. Each column represents a different sample. (B) The violin plot depicts the scaled count of *Il1rn* in myeloid subsets from both uninfected and infected samples. The scaled count were normalized using SCTransform method. IL-1Ra levels in the (C) skin and (D) serum of mice after 3 DPI of *C. auris* AR0387 or *C. albicans* SC5314 (*n =* 8-10 mice/group). Uninfected mice received 100 μL of 1X PBS on the day of infection. (E) Fungal burden in the skin tissue of indicated mice on 3 DPI of *C. auris* AR0387 or *C. albicans* SC5314 (*n =* 8-10 mice/group). Uninfected mice received 100 μL of 1X PBS on the day of infection. (F) The bar graph represents the neutrophil-killing activity of *C. auris* AR0387 in the presence and absence of recombinant IL-1Ra. (G) Skin fungal burden of the mice groups injected with neutralizing anti-mouse IL-1Ra monoclonal antibody or 100 µl of 1X PBS after 3 DPI of *C. auris* AR0387 in the murine skin. (*n =* 3-5 mice/group) (H-K) Quantification of IL-1Ra level and fungal burden in mice injected with anti-Ly6G antibody and Rat IgG2a isotype control (Isotype) for neutrophil depletion after 2 DPI of *C. auris* AR0387 (*n =* 6-8 mice/group). H) Representative flow plots and I) percentage and absolute number of CD11b^+^ Ly6G^+^ neutrophils in the infected skin tissue of mice injected with anti-Ly6G antibody or Rat IgG2a isotype control (Isotype) antibody. (J) IL-1Ra level and (K) CFU were determined from the homogenate of infected skin tissue. (L-O) Measurement of IL-1Ra production and fungal burden in clodrosome (C.L) and 1X PBS (Ctrl) injected mice groups for macrophage depletion after 2 DPI of *C. auris* AR0387 (*n =* 9-10 mice/group). (L) Representative flow plots of F4/80+ MHCII+ macrophage in the infected skin tissue of mice injected with clodrosome (C.L) or 1X PBS (Ctrl) injected mice groups. (M) IL-1Ra level and (N) CFU determined from the homogenate of infected skin tissue from 2 DPI of *C. auris* 0387 (n=12 mice/group).(O) IL-1Ra level and (P) CFU determined from the homogenate of infected skin tissue from 12 DPI of *C. auris* 0387 (n=8-10 mice/group). Error bars represent mean ± SEM. * p < 0.05, ** p <0.01, *** p <0.001, **** p <0.0001, NS, non-significant. Statistical significances were calculated using Mann–Whitney U. Abbreviations – DPI, days post-infection; C.L, clodrosome.

Our analysis revealed that *Il1rn* expression was highly expressed in neutrophils, followed by macrophages (Figure 5B). Given that *C. auris* persists in the skin long-term, the role of IL-1Ra in host defense against *C. auris* skin infection is unknown. Therefore, we investigated if the increased *Il1rn* identified in our scRNA analysis plays a role in host defense against *C. auris.* To confirm our scRNA data, we infected mice intradermally with *C. auris* and examined the IL-1Ra level by ELISA. In addition, mice intradermally infected with *C. albicans* was used to compare IL-1Ra levels with *C. auris*-infected groups. We identified that the skin tissue of *C. auris*-infected mice had significantly higher IL-1Ra levels compared with the uninfected and *C. albicans*-infected groups (Figure 5C). We found that all four *C. auris* clades (AR0381, AR0383, AR0285, and AR0387) induced comparable levels of IL-1Ra in skin tissues (Figure S6B and 6C). On the other hand, no significant difference was observed in the IL-1Ra level in the serum of uninfected, *C. auris,* and *C. albicans*-infected groups (Figure 5D). Furthermore, the increased level of IL-1Ra in the skin tissue of *C. auris*-infected mice corresponds to the increased fungal burden observed in *C. auris*-infected skin tissue compared with *C. albicans* infection groups (Figure 5E).

IL-1Ra binds with IL-1R [27, 30], and IL-1R signaling is critical for neutrophil function [26, 31, 32]. We investigated if the presence of IL-1Ra modulates the neutrophil killing of *C. auris.* The neutrophils isolated from the mouse peritoneum were assessed for *C. auris*-killing activity in the presence and absence of recombinant IL-1Ra. We identified that the neutrophils exhibited significantly decreased killing activity against *C. auris* in the presence of recombinant IL-1Ra (Figure 5F). Next, we examined the *in vivo* role of the secreted IL-1Ra in *C. auris* murine skin infection using neutralizing antibody. We found that *C. auris*-infected mice treated with the IL-1Ra neutralizing antibody exhibited significantly reduced fungal burdens in the skin tissue compared to untreated groups (Figure 5G). These findings indicate that the secreted IL-1Ra during *C. auris* skin infection decreases neutrophil function and potentially favors fungal growth in the skin tissue *in vivo*.

Next, we determined to examine the source of IL-1Ra during *C. auris* skin infection. Since neutrophils followed by macrophages highly expressed *Il1rn*, we examined if IL-1Ra produced by neutrophils and macrophages play a role in host defense against *C. auris* skin infection. We depleted neutrophils using an anti-Ly6G antibody, and as expected, the percentage and absolute number of CD11b^+^ Ly6G^+^ neutrophils were significantly decreased in the skin tissue of mice that received anti-Ly6G antibody compared to mice injected with isotype antibody (Figure 5H and 5I). However, surprisingly, we observed no significant difference in IL-1Ra levels in the skin tissue of mice that received anti-Ly6G antibody and isotype antibody (Figure 5J). On the other hand, fungal load was significantly increased in the skin tissue of mice that received anti-Ly6G antibody compared with mice that received isotype antibody (Figure 5K). These findings suggest neutrophils play a critical role in host defense against *C. auris* skin infection *in vivo*. Furthermore, depleting neutrophils alone did not considerably affect IL-1Ra levels in the skin tissue of mice infected with *C. auris*. Next, we depleted macrophages using clodronate liposomes to identify if macrophages are the source of IL-1Ra during *C. auris* skin infection. The percentage and absolute number of CD11b^+^ MHCII^+^ F4/80^+^ macrophages were significantly decreased in the skin tissue of mice that received clodronate liposomes compared to control groups (Figure 5L) (Figure S6D). Interestingly, we identified that IL-1Ra was significantly decreased in the skin tissue of mice after they received clodronate liposome compared with control groups (Figure 5M). Furthermore, the fungal load was significantly decreased in the skin tissue of mice that received clodronate liposome compared with control groups (Figure 5N). A difference of 1.6-fold in CFU/g of tissue was observed between the control and C.L. group. A similar trend was observed at a late time point, which was 12 DPI (Figure 5O and 5P). These findings suggest that macrophages are the major source of IL-1Ra during the early and late time points of *C. auris* skin infection. Next, we cultured the macrophages *in vitro*, stimulated with *C. auris,* and measured the IL-1Ra level in the culture supernatants after 16 hours. The IL-1Ra level in the culture supernatant of macrophages infected with *C. auris* was significantly higher than the uninfected macrophages. These results suggest that *C. auris* infection induces IL-1Ra production in the macrophage *ex vivo* (Figure S6E). To examine if the IL-1Ra produced by the macrophages can affect the neutrophil activity *ex vivo*, we collected the culture supernatant from the macrophages stimulated or unstimulated with *C. auris,* and this macrophage conditioned media (MΦ CM) was used to access the neutrophil killing activity of *C. auris*. The survival of *C. auris* in the neutrophil incubated with MΦ CM from infected macrophages was significantly increased compared to MΦ CM from uninfected macrophages (Figure S6F). This finding suggests that the macrophage-produced IL-1Ra limits the neutrophil-killing activity of *C. auris*. Collectively, our results indicate that the IL-1Ra produced by macrophages during *C. auris* skin infection decreases the neutrophil function, which is important for antifungal host defense against this emerging skin tropic fungal pathogen (Figure S7).

### *C. auris* uses a unique cell wall mannan layer to evade IL-1R signaling mediated host defense

IL-1R signaling is critical for neutrophil recruitment and function and plays an important role in the host defense against *C. albicans* in oral mucosa and systemic infection [31, 32]. However, the role of IL-1R signaling in host defense against *C. auris* skin infection is unknown. Therefore, we used mice lacking the IL-1 receptor (IL-1R) to investigate the IL-1R signaling in skin defense against *C. auris*. Surprisingly, we found that the percentage and absolute numbers of neutrophils in the *Il1r1-/-* mice infected with *C. auris* were not significantly affected and showed similar levels to wild-type (WT) infected groups after 3 DPI and 12 DPI (Figure 6A to 6C). We also observed no significant difference in the percentage and absolute numbers of macrophages in the *Il1r1-/-* mice after 3 DPI and 12 DPI (Figures 6D and 6E). Unlike *C. albicans* [31, 32], these findings indicate that neutrophil recruitment was not affected in *Il1r1-/-* mice infected with wild-type *C. auris* strain 0387. Since fungal PAMPs modulate the host response [12], and *C. auris* possesses a unique outer mannan layer compared to *C. albicans* [13], we hypothesized that the *C. auris* outer cell wall mannan layer controls IL-1R signaling. We used the CRISPR-CAS9 system to generate a *C. auris* mutant strain *(pmr1Δ)* that lacks an outer mannan layer as described previously[33]. The growth kinetics was unaltered in *C. auris pmr1Δ* [34]. WT and *Il1r1-/-* mice were infected with *pmr1Δ* strain to examine the neutrophil and macrophage recruitment. Interestingly, we found that neutrophil recruitment was significantly decreased in the *Il1r1-/-* mice infected with the *pmr1Δ* strain compared to wild-type (WT) mice after 3 DPI and 12 DPI (Figure 6F to 6H). No significant differences were observed in the recruitment of macrophages in the *Il1r1-/-* mice infected with the *pmr1Δ* strain (Figure 6I and 6J). These findings indicate that the unique mannan outer layer of *C. auris* controls IL-1R signaling-dependent neutrophil recruitment.

**Figure 6.**
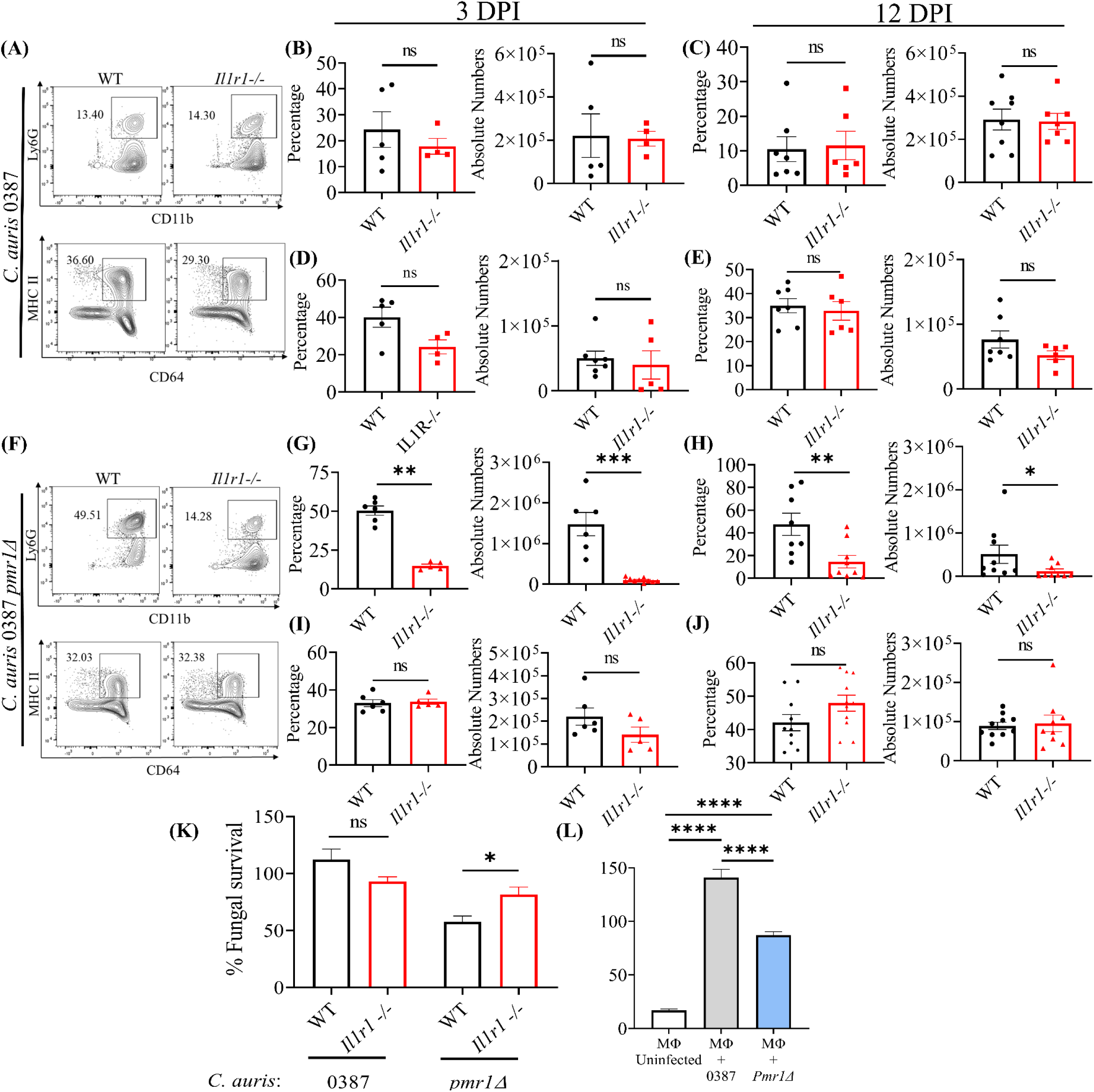
*C. auris* uses a unique cell wall mannan layer to evade IL-1R signaling mediated host defense. (A) The representative flow plots represents the percentage of CD11b^+^ Ly6G^+^ neutrophils and CD64^+^ MHCII^+^ macrophage in the WT and *IL-1R1^−/−^* mice skin tissue infected with *C. auris* 0387. The bar graph represents the percentage and absolute number of CD11b^+^ Ly6G^+^ neutrophils after (B) 3 DPI and (C) 14 DPI and the CD64^+^ MHCII^+^ macrophage after (D) 3 DPI and (E) 14 DPI in the WT and *IL-1R1^−/−^* mice skin tissue infected with *C. auris* 0387 (*n*=4-8 mice/group). (F) The representative flow plots represents the percentage of CD11b^+^ Ly6G^+^ neutrophils and CD64^+^ MHCII^+^ macrophage in the WT and *IL-1R1^−/−^* mice skin tissue infected with 0387 *pmr1Δ*. The bar graph represents the percentage and absolute number of CD11b^+^ Ly6G^+^ neutrophils after (G) 3 DPI and (H) 14 DPI and the CD64^+^ MHCII^+^ macrophage after (I) 3 DPI and (J) 14 DPI in the WT and *IL-1R1^−/−^*mice skin tissue infected with 0387 *pmr1Δ* (*n*=5-10 mice/group). For *C. auris* 0387 *pmr1Δ* infection 5 × 10^6^ CFU per mice were used. (K) The bar graph represents the fungal survival of *C. auris* 0387 and 0387 *pmr1Δ* primed with neutrophils isolated from bone marrow of WT and *IL-1R1^−/−^*mice. (*n*=6) (L) Measurement of IL-1Ra levels from the culture supernatant of BMDM uninfected or infected with live *C. auris* 0387 or 0387 *pmr1Δ* for 16 h. (*n* = 12). Error bars represent mean ± SEM. * p < 0.05, ** p <0.01, *** p <0.001, **** p <0.0001, NS, non-significant. Abbreviations - DPI, days post-infection; WT, wild type; BMDM, bone marrow-derived macrophages. Statistical significances were calculated using Mann–Whitney U.

Next, we investigated if *C. auris* mannan outer layer controls IL-1R signaling-dependent neutrophil function. Neutrophils isolated from WT and *Il1r1-/-* mice were infected with *C. auris* 0387 and 0387 *pmr1Δ* to determine fungal survival (Figure 6K). We identified no difference in fungal survival in neutrophils from WT and *Il1r1-/-* mice infected with the *C. auris* 0387 strain. However, we observed significantly higher fungal survival of the 0387 *pmr1Δ* strain in the *Il1r1-/-* neutrophils compared to the WT neutrophils (Figure 6K). These findings demonstrate that *C. auris*, through the outer mannan layer, evades IL-1R signaling mediated neutrophil killing. Since macrophages but not neutrophils are known to produce IL-1Ra during *C. albicans* systemic infection [35], we investigated whether the outer mannan layer controls macrophage IL-Ra production. BMDMs (bone marrow-derived macrophages) were infected with wild-type *C. auris* 0387 or 0387 *pmr1Δ* strain to examine IL-1Ra levels. We found a significant increase in the IL-Ra level in supernatants of macrophages infected with live wild-type *C. auris* 0387 strain compared to 0387 *pmr1Δ* strain, which lacks an outer mannan layer (Figure 6L). These findings suggest that outer mannan regulates IL-1Ra production. Taken together, our results revealed that through the unique mannan outer layer, *C. auris* evades IL-1R-signaling mediated neutrophil recruitment and killing to persist in the skin (Figure S7).

## Discussion

The single-cell transcriptomics platform is invaluable for capturing specific biological mechanisms and cellular heterogeneity among different cell populations in a tissue microenvironment. Given that *C. auris* possesses a unique cell wall, adhesin proteins, and colonizing factors to thrive in the skin [13, 17, 36], understanding the cell-type specific skin immune response to *C. auris* is critical to understanding the pathogenesis. In this study, comprehensive profiling of *C. auris* murine skin infection generated a single-cell transcriptome atlas encompassing all major immune cells and non-immune cells in the skin. Our scRNA dataset comprises ∼70,300 cells regulated during fungal infection of the skin tissue, revealing cell type-specific immune response during *C. auris* infection in the skin*. C. auris* can persist in the dermis for several months [37]. Therefore, we used the intradermal mouse model of infection that could potentially identify the host immune factors that regulate *C. auris* burden in skin tissue [20]. Our scRNA seq identified the accumulation of various innate and adaptive immune cells at the site of infection. In addition to phagocytic cells, DCs, and T cells, we identified that Th1 cells, NK cells, and non-immune cells such as fibroblast 3 cell type were highly accumulated in the skin tissue of *C. auris* infected mice. Recent studies suggest that IL-17-producing T cells are critical for skin defense against *C. auris* [4]. In this study, we observed that the depletion of neutrophils but not macrophages significantly increased the *C. auris* burden in skin tissue. These findings suggest neutrophils play a critical role in skin defense against *C. auris in vivo.* Future studies to understand the role of DCs, Th1 cells, NK cells, and fibroblasts in *C. auris* pathogenesis are important to understand the contribution of specific immune cells in skin defense against this emerging fungal pathogen.

Next, we examined the transcriptomic signature in individual cell types altered during *C. auris* skin infection *in vivo*. In myeloid cells, the cytokines and chemokines mediating the recruitment of phagocytic cells at the infection site and activation of antigen-presenting cells were highly upregulated during *C. auris* skin infection. The PRRs in the myeloid cells facilitate fungal recognition in the host [24, 38]. We observed the key PRRs, including TLRs, C-type lectin receptors, galectin receptors, and NLRs, were upregulated in myeloid cells, revealing the participation of selective immune receptors in host defense against *C. auris* during skin infection. Host AMPs such as *Lcn2, S100a8*, and *S100a9*, known to have direct antifungal activity, were highly upregulated in neutrophils. In addition, we identified the genes involved in nitric oxide metabolism and acute phase proteins were highly upregulated in different myeloid cells during *C. auris* skin infection. Our analysis of lymphoid cells revealed the expression of cytokines, chemokines, and TFs in CD4^+^ T cells, CD8^+^ T cells, and γδT cells during *C. auris* infection. In addition to the IL-17 pathway, which plays a critical role in antifungal host defense, including *C. auris* [4, 39], our findings suggest that *C. auris* infection highly upregulates the expression of *Ifng* in Th1 cells and NK cells. Previous findings indicate that IL-17 (Th17) but not IL-12 (Th1) is critical for host defense against *Candida albicans* in the skin tissue [40]. In addition to IL-17, our scRNA seq data highlights the potential involvement of IFNγ in *C. auris* infection. This could be partly due to the route of infection, as epicutaneous and intradermal infection elicits different immune responses [41, 42], and future studies are required to understand the functional importance of IFNγ (Th1) in *C. auris* skin infection. Non-immune cells, mainly fibroblast 1 and 3, showed increased expression of cytokines and chemokines. During *Staphylococcus aureus* skin infection, the dermal fibroblasts differentiate into adipocytes and produce AMP, mainly CAMP, for host defense [43]. Although CAMP was not highly expressed in fibroblasts during *C. auris* skin infection, our analysis identified the increased expression of other AMPs such as *Lcn2, Adm,* and *Ang* in fibroblast 1 and 3 cell types. The chemoattractants such as *Cxcl1, Cxcl2, Cxcl5, Cxcl12, Lcn2*, and *Il33* involved in recruiting immune cells such as neutrophils were highly expressed in various non-immune cells. These findings suggest that non-immune cells exhibit an active role in skin defense against *C. auris* through AMPs and by secreting soluble factors that assist in recruiting neutrophils to the site of infection. Collectively, our scRNA seq revealed the transcriptome changes in different immune and non-immune cell types during *C. auris* infection in mice skin tissue *in vivo* and identified the previously unknown immune and non-immune cell type-specific host responses against *C. auris* skin infection.

In the recent studies on the host transcriptome of *C. auris* in human peripheral blood mononuclear cells (PBMCs) and mouse BMDMs, the NOD-like receptor signaling pathway was only enriched in the *C. auris* infection but not in *C. albicans* infection [13, 14]. Similarly, in our scRNA seq dataset, the NOD-like receptor signaling pathway was significantly enriched in the myeloid subsets. Furthermore, the transcriptome of human PBMCs exposed *in vitro* with a purified manna layer of *C. auris* induces higher *Il1rn* expression compared to the *C. albicans* [13]. The *Il1rn* gene encoding IL-1Ra, which is significantly upregulated in different cell types identified in our scRNA seq data, was investigated for its role in host defense against *C. auris* skin infection. We identified that the depletion of macrophages, but not neutrophils, significantly decreased IL-1Ra concentrations in the skin tissue of mice infected with *C. auris,* suggesting the macrophage as the major source of IL-1Ra during *C. auris* skin infection. Furthermore, the neutrophil-killing activity of *C. auris* was significantly decreased in the presence of recombinant IL-1Ra, and mice treated with neutralizing antibodies significantly decreased the fungal load *in vivo.* A recent study on *C. albicans* using a systemic mouse model of infection revealed the role of secreted IL-1Ra by splenic macrophages limits the neutrophil activity in kidneys, contributing to the immune evasion of fungi [35]. We observed that the culture supernatant collected after *C. auris* stimulation of macrophage *in vitro* induced robust IL-1Ra production and limited the neutrophil-killing activity of *C. auris*. Our results, along with others [35], suggest that *C. albicans* and *C. auris* infection induces IL-1Ra in macrophages and decreases the killing activity of neutrophils to different host niches, including the kidney and skin, respectively. A recent study demonstrated that *C. auris* was less potent in activating IL-1β than *C. albicans,* evading macrophages through metabolic adaptation [44]. Furthermore, compared to *C. albicans,* we identified that *C. auris* infection induces significantly increased levels of IL-1Ra in the skin tissue. Although the intradermal infection of mice leads to the dissemination of fungi to kidneys and spleen [9], the systemic IL-1Ra level in serum in the mice infected with *C. auris* was unaltered. Previous findings suggest that IL-1Ra also favors the polarization of macrophages towards immunosuppressive M2 phenotype [45]. Macrophages play a protective role when a host encounters *Candida* pathogens. Our findings indicate the IL-1Ra-mediated immune evasion mechanisms employed by *C. auris* to modulate macrophages and persist in the skin. Furthermore, IL-1R signaling is critical for neutrophil recruitment and function and is important in the host defense against *C. albicans* [31, 32]. Surprisingly, our findings revealed that *C. auris*, through the unique mannan outer layer, evades IL-1R-mediated neutrophil recruitment and function in the skin. Collectively, our results demonstrate the IL-1Ra and IL-1R-mediated immune evasion mechanisms employed by *C. auris* to persist in the skin. However, IL-1Ra induction may differ between the murine and human in the skin, and intraspecies variation in the IL-1 regulation of innate immune response may exist [46]. Future studies using an *ex vivo* human skin infection model of *C. auris* might help to understand the IL-1Ra-mediated immune evasion. They may potentially correlate the findings from our mice studies to humans.

The overall conclusion is that in addition to known Th17-mediated antifungal defense mechanisms, our single-cell transcriptomics analysis identified Th1 cells, NK cells, and non-immune cells, such as fibroblast role in skin defense against *C. auris*. Our findings demonstrate that *C. auris* induces IL-1Ra in macrophages to decrease neutrophil function and evades IL-1R-mediated immune defense against *C. auris*. Taken together, our data from the comprehensive single-cell profiling of skin tissue revealed the previously unknown mechanisms by which *C. auris* modulates myeloid cells to persist in the skin. Furthermore, our unbiased transcriptomics identified the potential contribution of the IFNγ pathway in lymphoid cells and the involvement of non-immune cells in skin defense against *C. auris.* Findings from this study will form a strong platform for developing novel host and pathogen-directed antifungal therapeutics that potentially target IL-1Ra and fungal mannan biosynthesis, respectively. Furthermore, the host receptors and immune pathways identified in this study in individual skin cell types during *C. auris* infection *in vivo* will open the door for future studies to understand the pathogenesis of *C. auris* in the skin and develop novel therapeutics to prevent and treat this emerging skin tropic fungal pathogen.

## Author Contributions

ST designed the experiments and supervised the project. AB, ST, MN, VK, ML, DA, and JT performed data analysis. AD, DD, SG, AM, GB and ST carried out *in vivo* and *ex vivo* experiments. AB and ST wrote the original draft of the manuscript. All authors reviewed, edited, and approved the final version of the manuscript.

## Funding

This study was supported by NIAID (1R01AI177604 to ST) and the Division of Intramural Research of the NIAID, NIH (ZIA AI001175 to MSL).

### Experimental model and subject details

#### Mice

The C57BL/6J and *IL-1R1^−/−^* C57BL/6J strains of mice were bred and housed in pathogen-free conditions at the centrally managed animal facilities at Purdue University. C57BL/6 and *IL-1R1^−/−^* C57BL/6J strains were obtained from the Jackson Laboratory. Both old male and female mice of 6-8 weeks of age were used in the experiments. Animal studies and experimental protocols were approved by the Purdue University Institutional Animal Care and Use Committee (IACUC Protocol no. 2110002211).

#### Fungal culture

*Candida auris* AR0387, AR0381, AR0383, and AR0385 strains used in the study were procured from CDC AR Isolate Bank, USA, and *C. albicans* SC5314 strain was gifted from Dr. Andrew Koh, University of Texas Southwestern Medical Center, USA. Stock cultures of *C. auris, C. albicans* or *C. auris* 0387 *pmr1Δ* was stored at -80°C and subcultured on yeast peptone dextrose (YPD) agar plates at 37^0^C for 24 hours. Colonies from the YPD plates were cultured in YPD medium overnight at 37^0^ C, shaking at 250 rpm. The fungal cells were harvested and washed twice with sterile 1x PBS before murine infection. For ex *vitro* killing assays, *C. auris* was cultured in YPD at 30^0^ C for 24 hours. The fungal cells were washed with sterile 1x PBS and enumerated in a hemocytometer. The *C. auris* were preincubated with 10% mouse serum (Invitrogen, USA) at 37^0^C for 30 min. All the antibodies, ELISA kit, recombinant proteins, oligonucleotides, plasmids, and other reagents used in this study were provided in the (Table S6–S9).

## Method details

### *C. auris* murine skin infection

For scRNA seq, *C. auris* AR0387 was used for murine infection. Yeast cells were washed and resuspended in sterile 1x PBS before murine infection. The fungal cells were enumerated in a hemocytometer and diluted to 1-2 x 10^7^ yeast cells/ml. A six-to-eight-week-old C57BL/6J strain of mice was anesthetized, and dorsal skin hair was shaved as previously described [20]. Mice were injected intradermally with either 1-2 x 10^6^ yeast cells (*C. auris* infected group) or sterile 1× PBS (Uninfected group) using a 27 G 1’ hypodermic needle on the shaved area.

### Preparation of single-cell suspension from murine skin tissue for scRNA-seq

After 12 days post-infection (DPI), the mice were euthanized, and an approximately 80 – 120 mg weight of the dorsal skin tissue within 2 cm around the area of intradermal injection was collected in a 1.5 ml tube with digestion media (RPMI-1640 with 0.25-mg/mL Liberase TL and 1-µg/mL DNase). The tissue was transferred to a 6-well tissue culture plate with 5ml of digestion media and minced using ophthalmic scissors. The suspension is incubated for 1 hour 50 min at 37°C with 5% CO_2_ to ensure digestion of skin tissue. Then 1 mL of 0.25% trypsin-1-mM EDTA was added and incubated for another 10 min to separate the dermis from the epidermal surface as described previously [47]. After 2 hours of digestion, 4 ml of 1x PBS with 5% FBS was added to the suspension and pumped 8 times with a 10-mL syringe to mechanically dissociate the release of single cells from the tissues. Then the tissue clumps were removed by filtering with a 70 μm cell strainer, centrifuged at 300 x g for 7 min, and resuspended in RPMI-1640 with 10% FBS. Further, the fine tissue particles were removed by passing the single-cell suspension in a 40 μm cell strainer and a 30 μm cell strainer. The viability of the cells, presence of cell debris, and aggregating cells were assessed using trypan blue staining under a microscope. Two technical replicates for each sample were processed. Samples with more than 85% cell viability and less debris and aggregates were enumerated using a hemocytometer and adjusted to ∼10,000,000/mL for further Single-cell partition and RNA library preparation.

### Single-cell capturing, library construction, and 3’ RNA-sequencing

The Single-cell suspensions from the samples were diluted to 17,000 cells and loaded on a microfluidics chip Chromium Single Cell Instrument (10x Genomics) to target 10,000 cells. Single-cell GEMs (Gel Bead-In Emulsions) containing barcoded oligonucleotides and reverse transcriptase reagents were generated with the Next Gem single-cell reagent kit (10X Genomics). Following cell capture and cell lysis, cDNA was synthesized and amplified. At each step, the quality of cDNA and library was examined by a Bioanalyzer [48, 49]. The final libraries were sequenced on an Illumina NovaSeq 6000. 100-bp reads, including cell barcode and UMI sequences, and 100-bp RNA reads were generated.

### Bioinformatics analysis

The FASTQ files generated by Illumina sequencing were processed using the CellRanger (v7.1.0) pipeline from 10X Genomics, and the reads were mapped to mouse reference genome mm10-2020-A. A digital gene expression matrix was generated containing the raw UMI counts for each cell for each sample. Downstream analysis was performed using various functions in the Seurat package (v 4.0) [50–53]. Cells with fewer than 300 or more than 7500 unique feature counts or more than 20% mitochondrial reads were excluded. To address the ambient RNA contamination SoupX version 1.5.2[54] was utilized to identify and remove ambient RNA background noise from the data. After excluding low-quality cells, the count data was normalized using the SC Transform normalization workflow. Cell clusters were identified using ‘FindNeighbors’ and ‘FindClusters’ functions with the first 30 principal components at 0.6 resolution. We visualized our results in a two-dimensional space using UMAP. SingleR (v 2.0.0) package[55] was used for cell type annotation of the clusters using reference dataset GSE181720 [23]. Differential gene expression between infected and uninfected cells was performed using edgeR (v 3.38.4)[56]. Subclusters of lymphoid cell clusters identified above were identified by using the ‘resolution’ parameter from 0.1 to 1.0 in ‘FindClusters’ function to generate the detailed substructures of the clusters. We used pseudo-bulk analysis to combine single-cell RNA sequencing data by cell type and condition status, uninfected and infected. For each identified cell type within the seurat object, we extracted count matrices from both infected and uninfected samples. This extraction involved summing the UMIs (unique molecular identifiers) for each gene across all cells within a given cell type and condition, thereby creating pseudo-bulk samples representing each cell population’s collective gene expression profile under different experimental conditions. The differential expression analysis was performed using the edgeR package. We first normalized the data to account for differences in library sizes. Then we applied statistical models to identify genes that were significantly differentially expressed based on FDR ≤ 0.05 and Log2FC ≥ 2 or ≤ -2 between infected and uninfected conditions for each cell type. To visualize the results of our differential expression analysis, we generated volcano plots using the ggplot2 package in R. An ‘R’ package clusterProfiler (clusterProfiler_4.6.2)[57] was used to identify up-regulated KEGG pathways using differentially expressed genes between infected and uninfected cells when comparing different clusters and/or cell types.

### Depletion of neutrophils and macrophages in C57BL/6J mice

To deplete neutrophils in C57BL/6J mice, 200 µg of anti-mouse Ly-6G antibody (Biolegend, USA) were injected intraperitoneally on day -1 and +1 following *C. auris* skin infection. The control groups received isotype-matched control Rat IgG2a (Biolegend, USA) on the following days [58]. To deplete macrophages, C57BL/6J mice were injected with 1 mg of Clodrosome (Encapsula Nano Sciences) intraperitoneally on day -2 and +1 for 2 DPI, and alternative 3-4 days for 12 DPI following *C. auris* skin infection [59, 60]. PBS was injected into the control groups. Depletion of neutrophils and macrophages was assessed by flow cytometry. The skin fungal burden of the neutrophil or macrophage-depleted mice was determined from the infected skin tissue samples by homogenizing it in 1X PBS and plating in YPD supplemented with antibiotics, as described previously[20].

### Immune cell quantification by Flow Cytometry

Murine skin tissue from scRNA Seq validation and depletion experiments were subjected to immune cell quantification using flow cytometry. Cells were isolated from murine skin as described previously [20]. The single-cell suspension from the murine skin tissue was stained with LIVE/DEAD Fixable Yellow (Invitrogen, Waltham, MA, USA), followed by surface and intracellular markers for innate and adaptive panels as described previously [20]. For the depletion experiments, the macrophage population was determined by F4/80 antibody. Then the stained samples were acquired through Attune NxT Flow Cytometer (Invitrogen, Carlsbad, CA, USA) and analyzed using FlowJo software (Eugene, OR, USA).

### *In vivo* neutralization of IL-1Ra in the murine skin

To neutralize the IL-1Ra, 200 µg of neutralizing anti-mouse IL-1Ra monoclonal antibody were intradermally injected in the mice on day 0 (6 h before infection), +1 and +2 following *C. auris* skin infection [35, 61]. 1X PBS was given to the control groups on the following days. On 3 DPI the mice were euthanized, and skin fungal burden was determined by homogenizing the skin tissue and plating it in YPD plates supplemented with antibiotics as described previously[20].

### Blood sampling and serum separation

Peripheral blood was collected from uninfected and *C. albicans* and *C. auris*-infected mice on day 2 through the retro-orbital venous plexus route. Mice were anesthetized under isoflurane, and blood was collected using a heparinized microhematocrit capillary tube (Fisher Scientific, USA) in a microcentrifuge tube. Then, the blood was centrifuged at 500 × g for 10 min, and the separated serum was subjected to IL-1Ra ELISA.

### IL-1Ra quantification using enzyme-linked immunosorbent assays (ELISA)

IL-1Ra production in skin homogenate, culture supernatant, and serum was quantified using a mouse IL-1Ra/IL-1F3 DuoSet kit (RnD Systems) by following the manufacturer’s standard operating protocol. Skin tissue collected from mice was homogenized in PBS. The homogenates were centrifuged at 2500 × g for 10 min at 4^0^C, and the Halt protease inhibitor cocktail (ThermoFisher Scientific) was added to the supernatant and stored at -80^0^C. 20 µl of supernatant from skin homogenate, 50 µl of serum, 100 µl of BMDM culture supernatant were used for IL-1Ra ELISA.

### Isolation and culturing of murine primary neutrophils

Six to eight weeks old C57BL/6J or IL-1R1-/-mice were used for neutrophil isolation. As described previously, immune cells were isolated from mice’s peritoneal cavity [62]. Briefly, 7.5% of casein solution was injected intraperitoneally to stimulate immune cell infiltration. After 1 - 4 days, mice were euthanized, and the peritoneal cavity was washed with 1xPBS with 0.5% BSA, and the leukocytes were harvested from peritoneal lavage. Then the neutrophil was isolated from the buffy coat after centrifugation with Percoll at 1000 × g for 20 min as described previously [62]. For bone marrow-derived neutrophil isolation, the bone marrow cells were isolated from the femur by flushing the bone cavity with RPMI supplemented with 10% FBS and 2 mM EDTA. Then the RBC was lysed in 1.6% NaCl, and subjected to histopaque density gradient centrifugation for neutrophil separation as described previously[62].

### Construction of *pmr1Δ* deletion strain

The *PMR1* deletion in *C. auris* AR0387 was performed by CRISPR-mediated gene editing in *C. auris,* as described previously [33]. Briefly, the upstream and downstream regions of the *PMR1* gene were amplified and stretched with a 23 bp barcode, replacing the Open reading frame (ORF) to construct the repair template. Then, the gRNA cassette, CAS9 cassette, and the repair template were transformed into *C. auris* using lithium acetate, single-stranded carrier DNA, and polyethylene glycol method of competence induction. Then the transformed colonies were selected in the presence of NAT and screened for the *PMR1* deletion by colony PCR. The deletion of *PMR1* was confirmed using Sanger sequencing (Table S7).

### Isolation and culturing of murine bone marrow-derived macrophage (BMDM)

BMDM were isolated from 6–8 weeks old C57BL/6 mice as described previously[63]. Briefly, the bone marrow was harvested by flushing the bone cavity of the femur. Cells were cultured in DMEM supplemented with 10% FBS, 1× GlutaMAX, 1× Penicillin/Streptomycin, and 50 ng/mL M-CSF to induce macrophage differentiation as described[64]. After 5 -8 days, the adherent cells were separated, washed, and confirmed for CD11b^+^ MHCII^+^ by flow cytometry. 5 × 10^4^ cells/well of BMDM were cultured in a 96-well treated plate for 24 hours for efficient attachment. Then the macrophages were either stimulated at 1:1 MOI with live *C. auris* WT or *pmr1Δ* for 16 hours for the IL-1Ra ELISA. For preparation of macrophage conditioned media (MΦ CM), 2 × 10^5^ cells/well of BMDM were cultured in 48-well plate for 16 hours. The *C. auris* 0387 were stimulated with 1:1 MOI the *C. auris* MΦ CM, and the control conditioned media (ctrl MΦ CM) were unstimulated. Then 100 µl of supernatant was used for IL-1Ra level determination using ELISA.

### Neutrophil killing assay

To determine the *ex vivo* neutrophil killing of *C. auris* in the presence of IL-1Ra, 5 × 10^4^ neutrophils were seeded in a 96-well plate with RPMI 1640 media (ThermoFisher, USA) and pretreated with 100 ng/ml recombinant IL-1Ra for 30 min. Further, *C. auris* AR0387 was incubated with 10% mouse serum for 30 min and transferred to the 96-well plate at MOI of 1: 0.25. The plates were incubated at 37^0^C for 3 hours. Then, the cells were lysed by resuspending in 0.02% triton X-100 for 5 min, and the lysate was serially diluted and plated in YPD agar with antibiotics. The colonies were enumerated after incubating plates at 37^0^C for 24 hours to calculate the neutrophil-killing activity. For fungal survival assay, *C. auris* 0387 or 0387 *pmr1Δ* were primed with 5 × 10^4^ bone marrow neutrophils isolated from WT or *IL-1R1-/-* mice at 1: 2 MOI and the percentage of fungal survival were determined by plated in YPD agar with antibiotics [65, 66]. For fungal survival in macrophage conditioned media (MΦ CM), the neutrophil and *C. auris* were primed at 1:0.25 MOI in presence of MΦ CM at various concentrations and for 50 % MΦ CM, the MΦ CM were diluted in DMEM supplemented with 10% FBS. The percentage of fungal survival were determined by plated in YPD agar with antibiotics.

## References

1. Ahmad, S. and W. Alfouzan, Candida auris: Epidemiology, Diagnosis, Pathogenesis, Antifungal Susceptibility, and Infection Control Measures to Combat the Spread of Infections in Healthcare Facilities. Microorganisms, 2021. 9(4).

2. WHO fungal priority pathogens list to guide research, development and public health action, Geneva: World Health Organization; 2022 Licence: CC BY-NC-SA 30 IGO, https://www.who.int/publications/i/item/9789240060241. 2022, WHO.

3. Kadri, S.S., Key Takeaways From the U.S. CDC’s 2019 Antibiotic Resistance Threats Report for Frontline Providers. Crit Care Med, 2020. 48(7): p. 939–945.

4. Huang, X., et al., Murine model of colonization with fungal pathogen Candida auris to explore skin tropism, host risk factors and therapeutic strategies. Cell Host Microbe, 2021. 29(2): p. 210–221 e6.

5. Shastri, P.S., et al., Candida auris candidaemia in an intensive care unit - Prospective observational study to evaluate epidemiology, risk factors, and outcome. J Crit Care, 2020. 57: p. 42–48.

6. Proctor, D.M., et al., One population, multiple lifestyles: Commensalism and pathogenesis in the human mycobiome. Cell Host Microbe, 2023. 31(4): p. 539–553.

7. Sparber, F., et al., The Skin Commensal Yeast Malassezia Triggers a Type 17 Response that Coordinates Anti-fungal Immunity and Exacerbates Skin Inflammation. Cell Host Microbe, 2019. 25(3): p. 389–403 e6.

8. Proctor, D.M., et al., Integrated genomic, epidemiologic investigation of Candida auris skin colonization in a skilled nursing facility. Nature Medicine, 2021. 27(8): p. 1401–1409.

9. Towns, K.A., A. Datta, and S. Thangamani, Intradermal infection and dissemination of Candida auris in immunocompetent and immunocompromised mouse models. Microbiol Spectr, 2024: p. e0012724.

10. Pekmezovic, M., et al., Candida pathogens induce protective mitochondria-associated type I interferon signalling and a damage-driven response in vaginal epithelial cells. Nat Microbiol, 2021. 6(5): p. 643–657.

11. Lionakis, M.S., R.A. Drummond, and T.M. Hohl, Immune responses to human fungal pathogens and therapeutic prospects. Nat Rev Immunol, 2023. 23(7): p. 433–452.

12. Briard, B., et al., Fungal cell wall components modulate our immune system. Cell Surf, 2021. 7: p. 100067.

13. Bruno, M., et al., Transcriptional and functional insights into the host immune response against the emerging fungal pathogen Candida auris. Nat Microbiol, 2020. 5(12): p. 1516–1531.

14. Wang, Y., et al., Innate immune responses against the fungal pathogen Candida auris. Nat Commun, 2022. 13(1): p. 3553.

15. Yan, L., et al., Unique Cell Surface Mannan of Yeast Pathogen Candida auris with Selective Binding to IgG. ACS Infect Dis, 2020. 6(5): p. 1018–1031.

16. Seiser, S., et al., Native human and mouse skin infection models to study Candida auris-host interactions. Microbes Infect, 2024. 26(1-2): p. 105234.

17. Santana, D.J., et al., A Candida auris-specific adhesin, Scf1, governs surface association, colonization, and virulence. Science, 2023. 381(6665): p. 1461–1467.

18. Herrada, J., et al., In Vivo Skin Colonization and Decolonization Models for Candida auris. Methods Mol Biol, 2022. 2517: p. 269–285.

19. Bryak, G., et al., Yeast and filamentous Candida auris stimulate distinct immune responses in the skin. mSphere, 2024: p. e0005524.

20. Datta, A., et al., Differential skin immune responses in mice intradermally infected with Candida auris and Candida albicans. Microbiol Spectr, 2023: p. e0221523.

21. Solá, P., et al., Targeting lymphoid-derived IL-17 signaling to delay skin aging. Nature Aging, 2023. 3(6): p. 688–704.

22. Cui, A., et al., Dictionary of immune responses to cytokines at single-cell resolution. Nature, 2024. 625(7994): p. 377–384.

23. Venugopal, G., et al., In vivo transcriptional analysis of mice infected with Leishmania major unveils cellular heterogeneity and altered transcriptomic profiling at single-cell resolution. PLOS Neglected Tropical Diseases, 2022. 16(7): p. e0010518.

24. Netea, M.G., et al., Immune defence against Candida fungal infections. Nature Reviews Immunology, 2015. 15(10): p. 630–642.

25. Blake, J.A., et al., Mouse Genome Database (MGD): Knowledgebase for mouse–human comparative biology. Nucleic Acids Research, 2020. 49(D1): p. D981–D987.

26. Mantovani, A., et al., Interleukin-1 and Related Cytokines in the Regulation of Inflammation and Immunity. Immunity, 2019. 50(4): p. 778–795.

27. Carter, D.B., et al., Purification, cloning, expression and biological characterization of an interleukin-1 receptor antagonist protein. Nature, 1990. 344(6267): p. 633–638.

28. Martin, P., et al., Intracellular IL-1 Receptor Antagonist Isoform 1 Released from Keratinocytes upon Cell Death Acts as an Inhibitor for the Alarmin IL-1α. The Journal of Immunology, 2020. 204(4): p. 967–979.

29. Kanangat, S., et al., Novel Functions of Intracellular IL-1ra in Human Dermal Fibroblasts: Implications in the Pathogenesis of Fibrosis. Journal of Investigative Dermatology, 2006. 126(4): p. 756–765.

30. Hannum, C.H., et al., Interleukin-1 receptor antagonist activity of a human interleukin-1 inhibitor. Nature, 1990. 343(6256): p. 336–340.

31. Altmeier, S., et al., IL-1 Coordinates the Neutrophil Response to C. albicans in the Oral Mucosa. PLoS Pathog, 2016. 12(9): p. e1005882.

32. Bellocchio, S., et al., The contribution of the Toll-like/IL-1 receptor superfamily to innate and adaptive immunity to fungal pathogens in vivo. J Immunol, 2004. 172(5): p. 3059–69.

33. Ennis, C.L., A.D. Hernday, and C.J. Nobile, A Markerless CRISPR-Mediated System for Genome Editing in Candida auris Reveals a Conserved Role for Cas5 in the Caspofungin Response. Microbiol Spectr, 2021. 9(3): p. e0182021.

34. Horton, M.V., et al., Candida auris Cell Wall Mannosylation Contributes to Neutrophil Evasion through Pathways Divergent from Candida albicans and Candida glabrata. mSphere, 2021. 6(3): p. e0040621.

35. Gander-Bui, H.T.T., et al., Targeted removal of macrophage-secreted interleukin-1 receptor antagonist protects against lethal Candida albicans sepsis. Immunity, 2023. 56(8): p. 1743–1760 e9.

36. Balakumar, A., D. Bernstein, and S. Thangamani, The adhesin SCF1 mediates Candida auris colonization. Trends in Microbiology, 2024. 32(1): p. 4–5.

37. Huang, X., et al., Murine model of colonization with fungal pathogen Candida auris to explore skin tropism, host risk factors and therapeutic strategies. Cell Host & Microbe, 2021. 29(2): p. 210–221.e6.

38. Netea, M.G., et al., An integrated model of the recognition of Candida albicans by the innate immune system. Nature Reviews Microbiology, 2008. 6(1): p. 67–78.

39. Conti, H.R. and S.L. Gaffen, IL-17–Mediated Immunity to the Opportunistic Fungal Pathogen Candida albicans. The Journal of Immunology, 2015. 195(3): p. 780–788.

40. Kagami, S., et al., IL-23 and IL-17A, but not IL-12 and IL-22, are required for optimal skin host defense against Candida albicans. J Immunol, 2010. 185(9): p. 5453–62.

41. Linehan, J.L., et al., Non-classical Immunity Controls Microbiota Impact on Skin Immunity and Tissue Repair. Cell, 2018. 172(4): p. 784–796.e18.

42. Kashem, S.W., et al., Candida albicansMorphology and Dendritic Cell Subsets Determine T Helper Cell Differentiation. Immunity, 2015. 42(2): p. 356–366.

43. Zhang, L.-j., et al., Dermal adipocytes protect against invasive Staphylococcus aureus skin infection. Science, 2015. 347(6217): p. 67–71.

44. Weerasinghe, H., et al., Candida auris uses metabolic strategies to escape and kill macrophages while avoiding robust activation of the NLRP3 inflammasome response. Cell Rep, 2023. 42(5): p. 112522.

45. Luz-Crawford, P., et al., Mesenchymal Stem Cell-Derived Interleukin 1 Receptor Antagonist Promotes Macrophage Polarization and Inhibits B Cell Differentiation. Stem Cells, 2016. 34(2): p. 483–92.

46. Tahtinen, S., et al., IL-1 and IL-1ra are key regulators of the inflammatory response to RNA vaccines. Nature Immunology, 2022. 23(4): p. 532–542.

47. Sakamoto, K., et al., Flow cytometry analysis of the subpopulations of mouse keratinocytes and skin immune cells. STAR Protocols, 2022. 3(1): p. 101052.

48. Macosko, Evan Z., et al., Highly Parallel Genome-wide Expression Profiling of Individual Cells Using Nanoliter Droplets. Cell, 2015. 161(5): p. 1202–1214.

49. Zheng, G.X.Y., et al., Massively parallel digital transcriptional profiling of single cells. Nature Communications, 2017. 8(1): p. 14049.

50. Stuart, T., et al., Comprehensive Integration of Single-Cell Data. Cell, 2019. 177(7): p. 1888–1902.e21.

51. Satija, R., et al., Spatial reconstruction of single-cell gene expression data. Nature Biotechnology, 2015. 33(5): p. 495–502.

52. Butler, A., et al., Integrating single-cell transcriptomic data across different conditions, technologies, and species. Nature Biotechnology, 2018. 36(5): p. 411–420.

53. Hao, Y., et al., Integrated analysis of multimodal single-cell data. Cell, 2021. 184(13): p. 3573–3587.e29.

54. Young, M.D. and S. Behjati, SoupX removes ambient RNA contamination from droplet-based single-cell RNA sequencing data. GigaScience, 2020. 9(12): p. giaa151.

55. Aran, D., et al., Reference-based analysis of lung single-cell sequencing reveals a transitional profibrotic macrophage. Nature Immunology, 2019. 20(2): p. 163–172.

56. Robinson, M.D., D.J. McCarthy, and G.K. Smyth, edgeR: a Bioconductor package for differential expression analysis of digital gene expression data. Bioinformatics, 2010. 26(1): p. 139–140.

57. Yu, G., et al., clusterProfiler: an R Package for Comparing Biological Themes Among Gene Clusters. OMICS: A Journal of Integrative Biology, 2012. 16(5): p. 284–287.

58. Carr, K.D., et al., Specific depletion reveals a novel role for neutrophil-mediated protection in the liver during Listeria monocytogenes infection. European Journal of Immunology, 2011. 41(9): p. 2666–2676.

59. Wang, J. and P. Kubes, A Reservoir of Mature Cavity Macrophages that Can Rapidly Invade Visceral Organs to Affect Tissue Repair. Cell, 2016. 165(3): p. 668–678.

60. Hiengrach, P., et al., Macrophage depletion alters bacterial gut microbiota partly through fungal overgrowth in feces that worsens cecal ligation and puncture sepsis mice. Scientific Reports, 2022. 12(1): p. 9345.

61. Tanaka, R., et al., Efficient drug delivery to lymph nodes by intradermal administration and enhancement of anti-tumor effects of immune checkpoint inhibitors. Cancer Treatment and Research Communications, 2023. 36: p. 100740.

62. Swamydas, M., et al., Isolation of Mouse Neutrophils. Curr Protoc Immunol, 2015. **110**: p. 3.20.1–3.20.15.

63. Zhang, X., R. Goncalves, and D.M. Mosser, The isolation and characterization of murine macrophages. Curr Protoc Immunol, 2008. **Chapter 14**: p. 14 1 1–14 1 14.

64. Haag, S.M. and A. Murthy, Murine Monocyte and Macrophage Culture. Bio Protoc, 2021. 11(6): p. e3928.

65. Johnson Chad, J., et al., Emerging Fungal Pathogen Candida auris Evades Neutrophil Attack. mBio, 2018. 9(4): p. 10.1128/mbio.01403-18.

66. Wu, S.-Y., et al., Candida albicans triggers NADPH oxidase-independent neutrophil extracellular traps through dectin-2. PLOS Pathogens, 2019. 15(11): p. e1008096.

