## Supplementary Materials for "Single-Cell Transcriptomics Unveils Skin Cell Specific Antifungal Immune Responses and IL-1Ra-IL-1R Immune Evasion Strategies of Emerging Fungal Pathogen *Candida auris*"

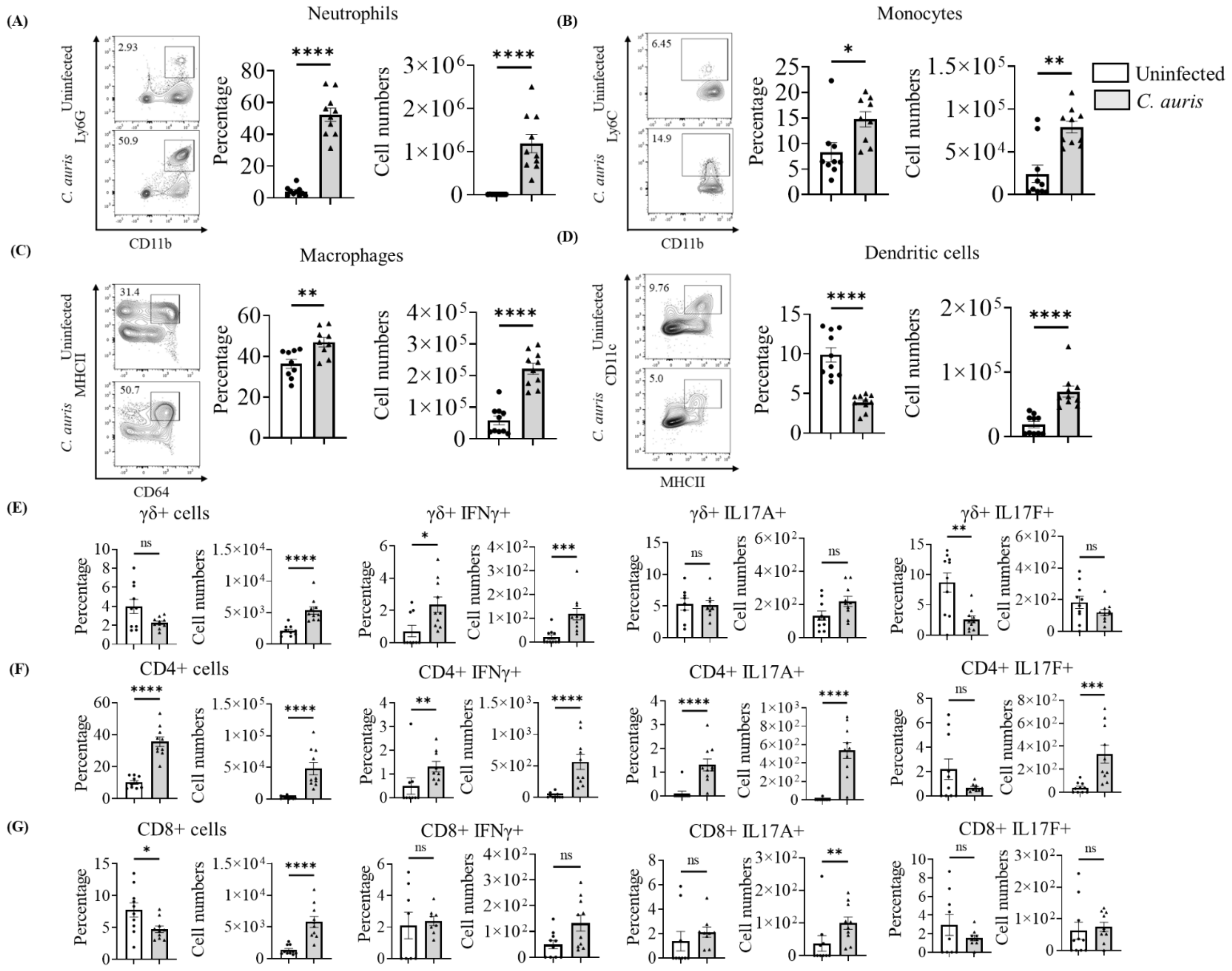


**Figure S1:** Flow cytometric quantification of major immune cells from the mice skin tissue of the infected and uninfected groups after 12-days-post-infection. The representative flow plot and the bar graph represent the percentage and absolute number of (A) neutrophils, (B) Monocytes, (C) Macrophages, and (D) dendritic cells in the *C. auris* infected group compared to the uninfected group. The bar graph represents the percentage and absolute number of (E) γδ+ cells, γδ+ IFNγ+ cells, γδ+ IL17A+ cells, and γδ+ IL17F+ cells, (F) CD4+ cells, CD4+ IFNγ+ cells, CD4+ IL17A+ cells, CD4+ and IL17F+ cells, (G) CD8+ cells, CD8+ IFNγ+ cells, CD8+ IL17A+ cells, CD8+ and IL17F+ cells. Twelve mice were used from each group, and the error bar represents the mean ± SEM. * $p < 0.05$, ** $p < 0.01$, *** $p < 0.001$, **** $p < 0.0001$.

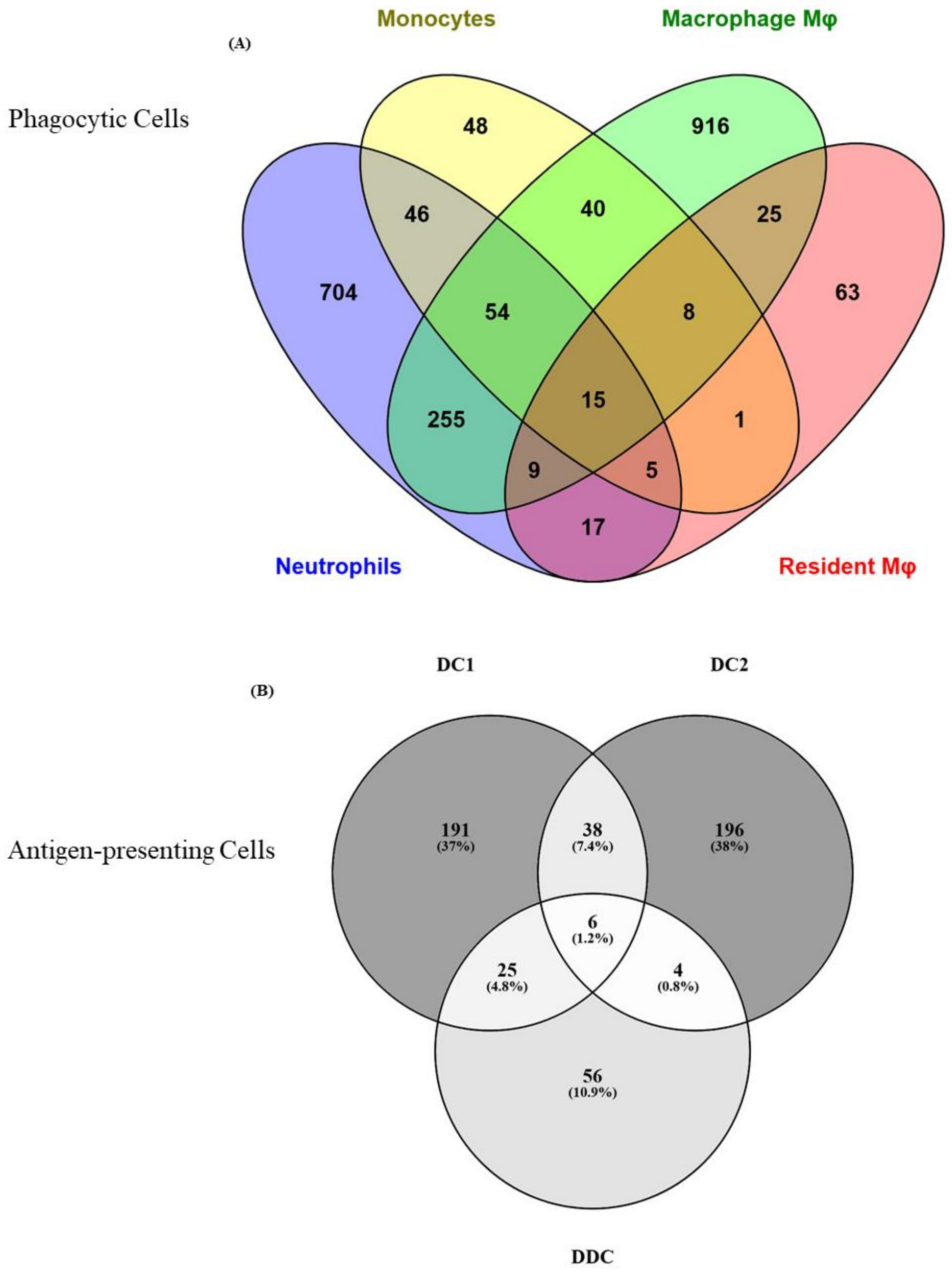


**Figure S2:** Venn diagram showing the number of upregulated genes shared within the (A) phagocytic cells and (B) antigen presenting cells upon *C. auris* infection.

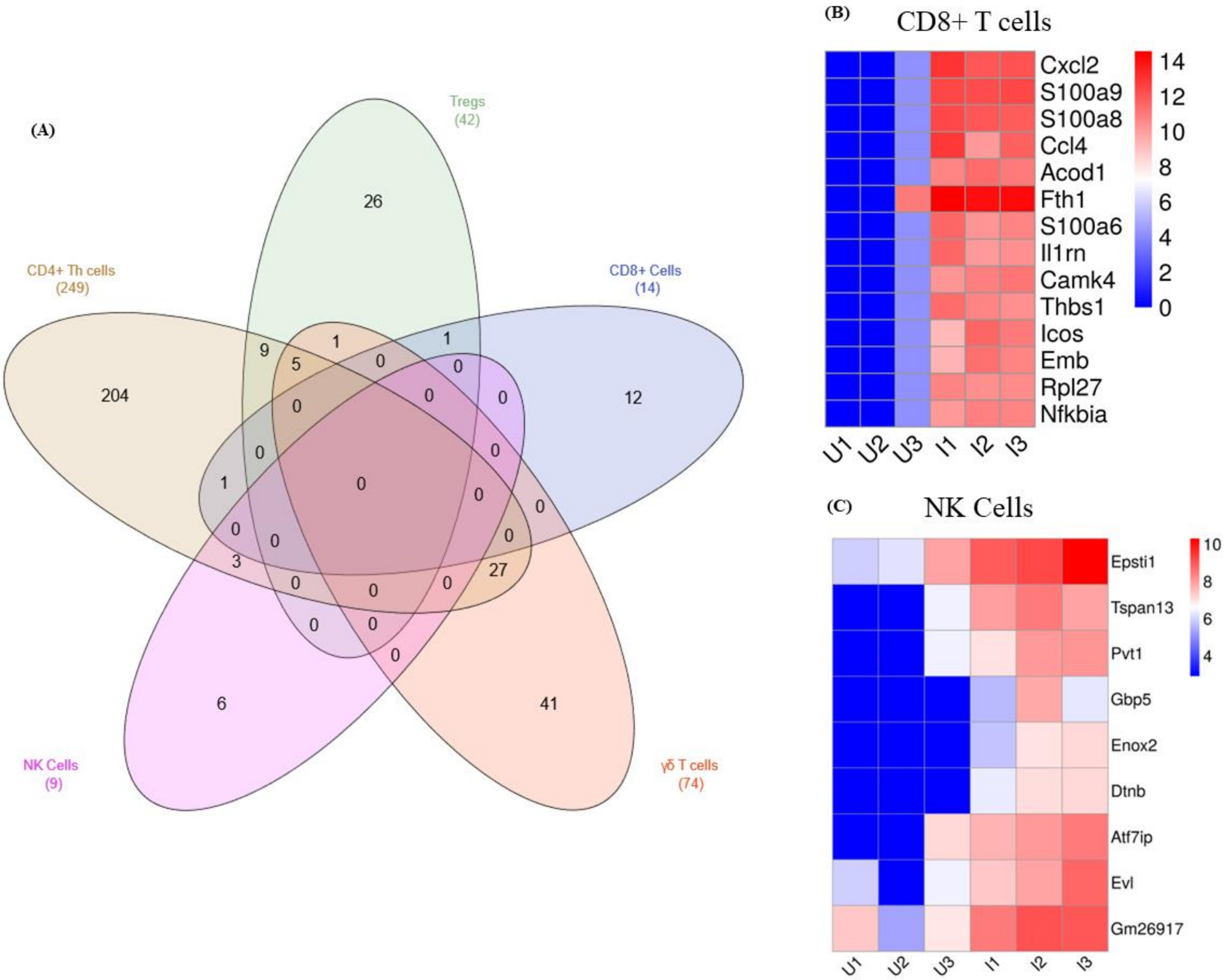


**Figure S3:** (A) Venn diagram showing the number of upregulated genes shared within lymphoid subsets. The heatmap represents the expression of the significantly upregulated genes in (B) CD8+ and (C) NK cells in uninfected and infected groups. Upregulated genes with Log 2-fold change + 2 and FDR > 5% were represented. The normalized gene counts were plotted in the heatmap, and the scale indicates red for high, blue for low, and white for moderate expression in the samples. Each column represents a different sample.

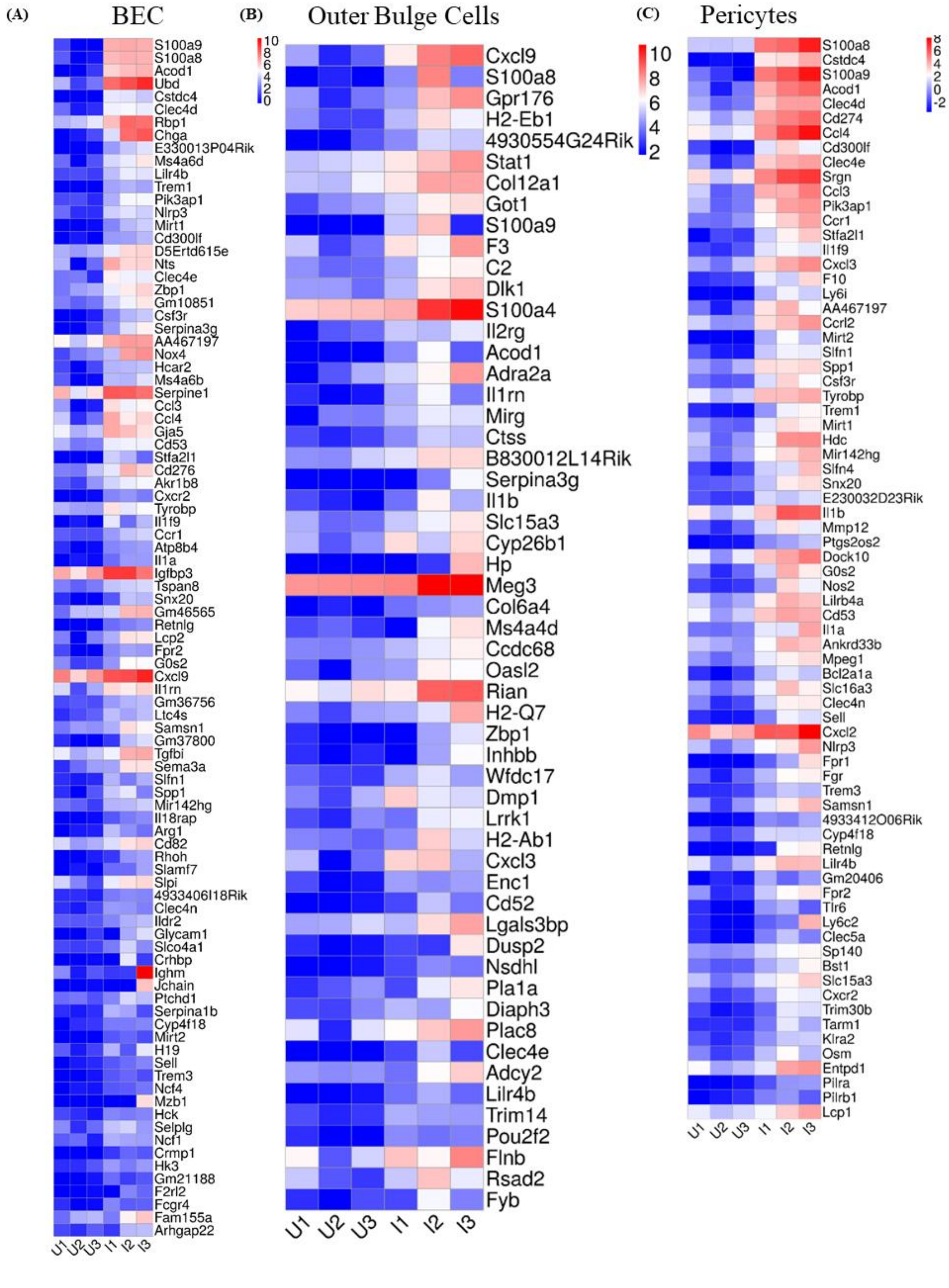


**Figure S4:** The heatmap represents the expression of the significantly upregulated genes in (A) BEC (B) outer bulge cells and (C) pericytes cells in uninfected and infected groups. Upregulated genes with Log 2-fold change ≥ 2 and FDR > 5% were represented. The normalized gene counts were plotted in the heatmap, and the scale indicates red for high, blue for low, and white for moderate expression in the samples. Each column represents a different sample.

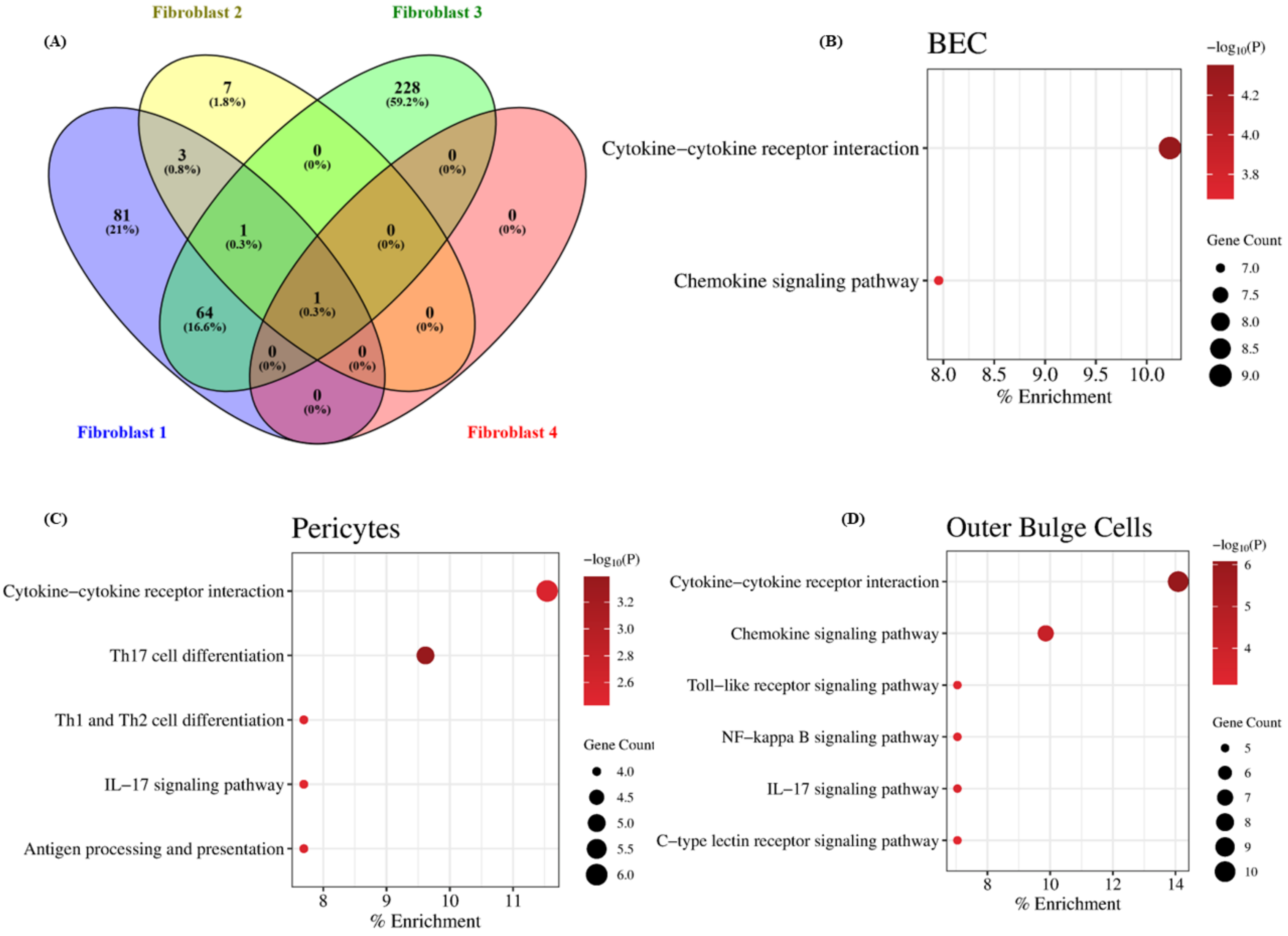


**Figure S5:** (A) Venn diagram showing the number of significantly upregulated genes shared within the identified fibroblast subsets. The bubble plot represents the KEGG pathways of the enriched upregulated genes of the (B) BEC, (C) pericytes and (D) outer bulge cells. The X-axis denotes the percentage enrichment of the KEGG pathways.

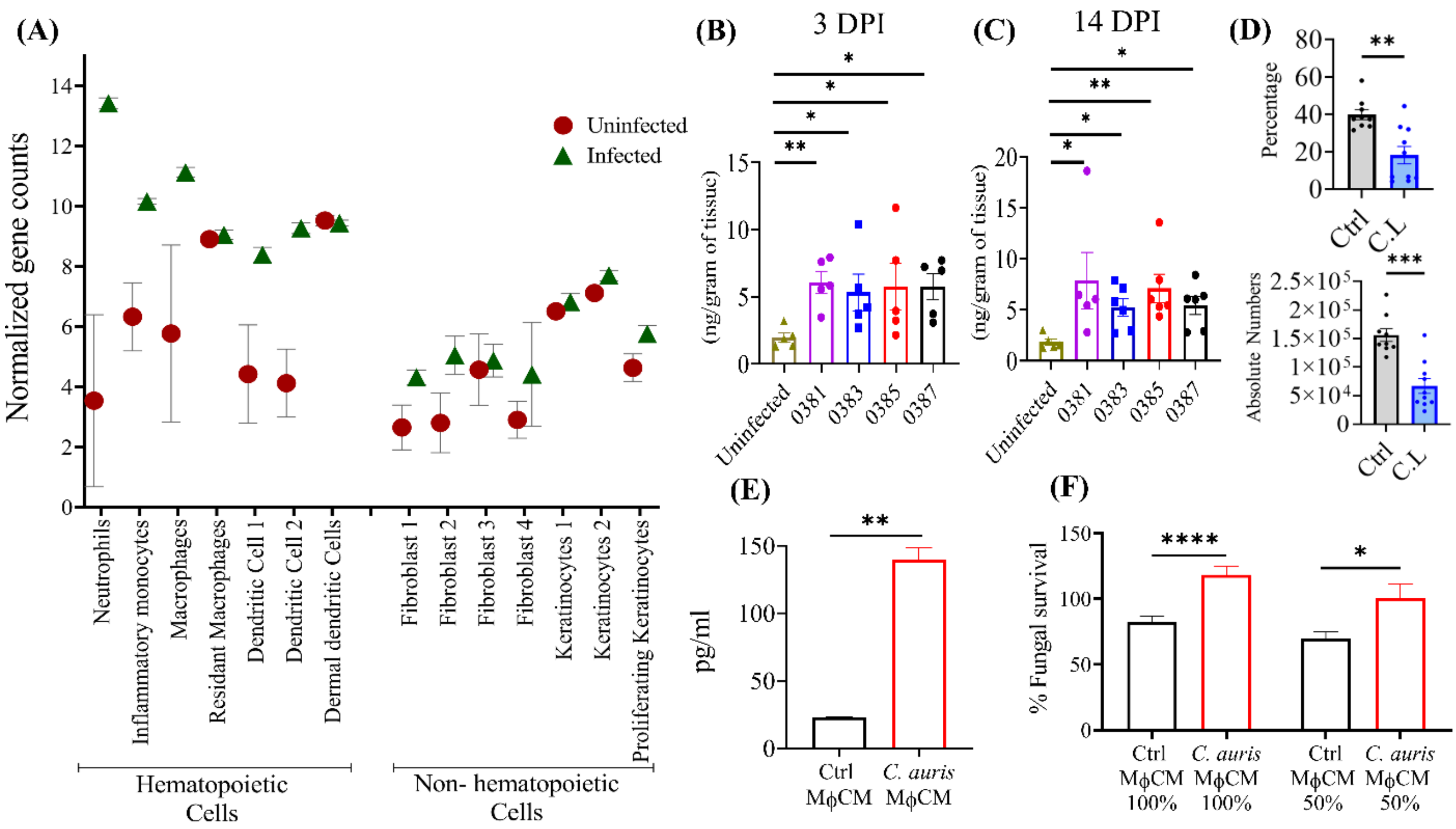


**Figure S6:** (A) The normalized read counts of *Il1rn* gene in the hematopoietic and non-hematopoietic cell types from the scRNA-seq dataset. Both uninfected and infected samples were plotted and the mean were represented with the error bar. The IL-1Ra level in skin tissues of mice groups received PBS, *Candida auris* South Asian clade AR0387, East Asian clade AR0381, African clade AR0383, or South American clade AR0385 after day. (B) 3 p.i. and (C) 14 p.i. (*n* = 5-7 mice/group). (D) Percentage and absolute number of F4/80$^+$ MHCII$^+$ macrophage in the infected skin tissue of mice injected with clodrosome (C.L) or 1X PBS (Ctrl) injected mice groups. (E) Measurement IL-1Ra levels from the culture supernatant of BMDM alone or BMDM stimulated with *C. auris* 0387 for 16 h. (*n* = 12). Error bars represent mean ± SEM. ** p <0.01. (F) The bar graph represents the fungal survival of *C. auris* 0387 primed with neutrophils in the presence of MΦ CM collected from BMDM alone (Ctrl MΦ CM) or BMDM stimulated with *C. auris* 0387 (*C. auris* MΦ CM) for 16 h (*n* = 16 to 19). 50% CM was diluted with complete DMEM. Error bars represent mean ± SEM. * p <0.05, **** p <0.0001. Abbreviations - BMDM, bone marrow-derived macrophages; MΦ CM, macrophage conditioned medium. Statistical significances were calculated using Mann–Whitney U.

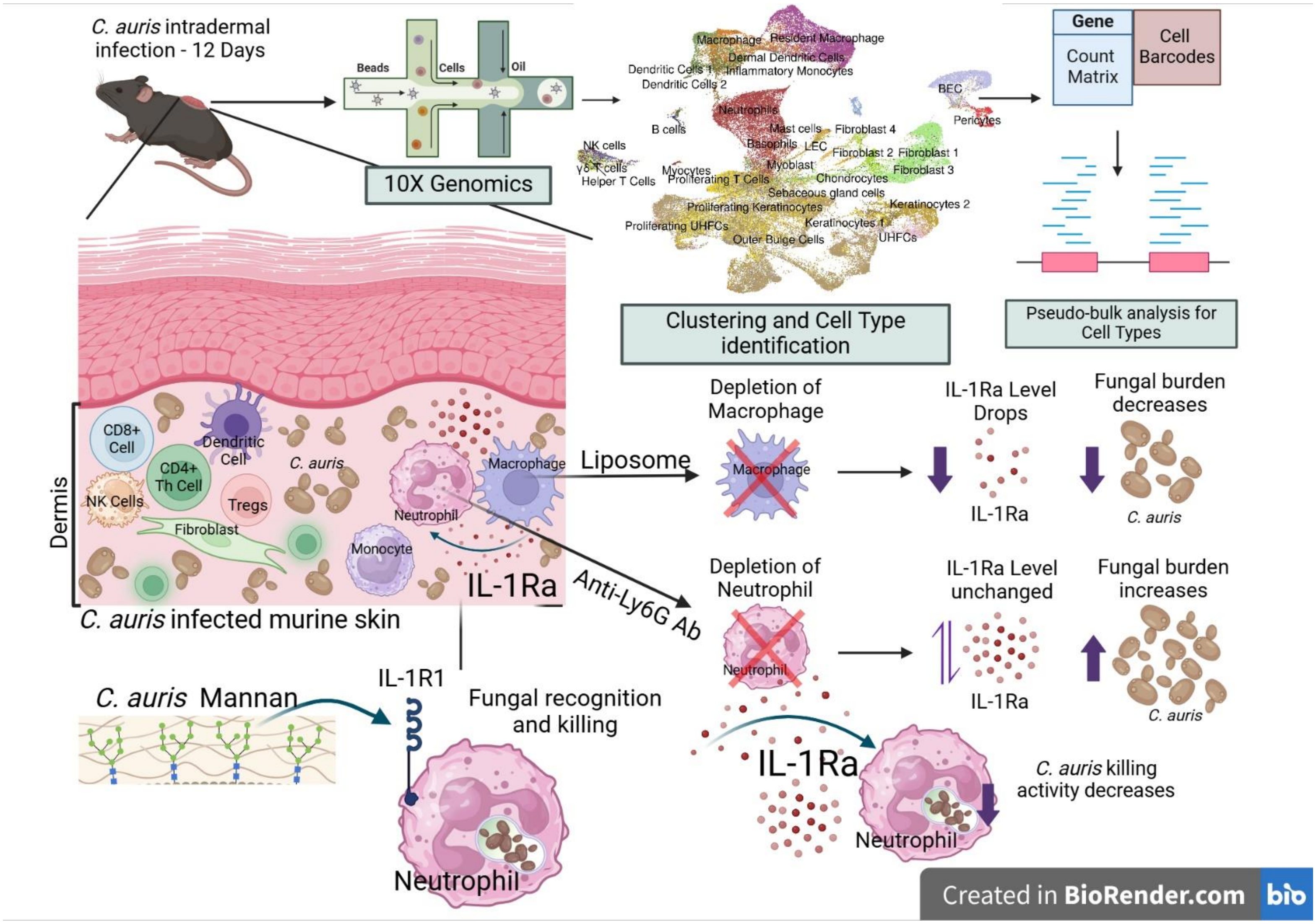


**Figure S7:** The illustration of *in vivo* scRNA seq of murine skin infected with *C. auris* revealing IL-1Ra mediated host evasion mechanism for fungal survival in the skin.

**Table S1:** The number of cells in each cell type identified in the infected and uninfected groups. The mean cell numbers of the cell type in the uninfected and infected samples were represented. The fold changes in the cell types recruited in the infected samples compared to the uninfected control were represented.

| Cell Types | No. of Cells in Uninfected | No. of Cells in Infected | Fold Change in Cell Type recruited |
|---|---|---|---|
| B cells | 55 | 101 | 0.836363636 |
| BEC | 2083 | 804 | -0.614018243 |
| Basophils | 8 | 33 | 3.125 |
| Chondrocytes | 109 | 9 | -0.917431193 |
| Cytotoxic T cells | 91 | 387 | 3.252747253 |
| DN T-lymphocytes | 8 | 3 | -0.625 |
| Dendritic Cells 1 | 129 | 643 | 3.984496124 |
| Dendritic Cells 2 | 24 | 240 | 9 |
| Dermal Dendritic Cells | 1197 | 1284 | 0.072681704 |
| Fibroblast 1 | 4486 | 984 | -0.780650914 |
| Fibroblast 2 | 999 | 162 | -0.837837838 |
| Fibroblast 3 | 49 | 355 | 6.244897959 |
| Fibroblast 4 | 390 | 40 | -0.897435897 |
| Helper T Cells | 32 | 850 | 25.5625 |
| Inflammatory Monocytes | 57 | 758 | 12.29824561 |
| Keratinocytes 1 | 1650 | 351 | -0.787272727 |
| Keratinocytes 2 | 2313 | 381 | -0.835278859 |
| LEC | 204 | 72 | -0.647058824 |
| Macrophage | 12 | 3050 | 253.1666667 |
| Mast cells | 63 | 34 | -0.46031746 |
| Myoblast | 1205 | 250 | -0.79253112 |
| Myocytes | 96 | 45 | -0.53125 |
| NK cells | 24 | 239 | 8.958333333 |
| Neutrophils | 8 | 7548 | 942.5 |
| Outer Bulge Cells | 13249 | 4292 | -0.676051023 |
| Pericytes | 556 | 214 | -0.615107914 |
| Proliferating Keratinocytes | 4438 | 1238 | -0.721045516 |
| Proliferating T Cells | 71 | 141 | 0.985915493 |
| Proliferating UHFCs | 1206 | 406 | -0.663349917 |
| Resident Macrophage | 2778 | 3975 | 0.430885529 |
| Schwann cells | 363 | 98 | -0.730027548 |
| Sebaceous gland cells | 1031 | 374 | -0.637245393 |
| Tregs | 45 | 254 | 4.644444444 |

**Table S2:** The total UMIs and annotated genes identified in the infected and uninfected groups. The mean UMIs in uninfected and infected sample are represented in the table.

| Groups | No. of samples | Mean UMIs | Mean genes |
|---|---|---|---|
| Uninfected | 3 | 91294458 | 465724 |
| Infected | 3 | 64566434 | 429009 |

**Table S3:** The DEGs of myeloid subsets enriched in the KEGG pathways upon *C. auris* murine skin infection.

| KEGG Pathways | DEGs enriched in the pathway |
|---|---|
| Phagosome | **Neutrophils** - *Ctss, Sec61g, H2-T23, Canx, Itgav, Fcgr3, Atp6v0d2, Atp6v1c1, Atp6v1b2, H2-Q10, Ncf1, Atp6v1g1, Itgb1, Cd14, H2-K1, Atp6v0c, Rab7, Atp6v1a, H2-Q6, M6pr, Atp6v1d, Tap2, Olr1, Rac1, Tap1, Lamp2, Tubb6, Lamp1, Tlr6, Tlr4, Stx7, Tlr2, C3, H2-Q7, H2-T22, H2-Aa, H2-Ab1, Cybb, Atp6v1e1, Atp6v0d1, Atp6v0a1, H2-D1, Eea1* and *Atp6v0e*<br>**Inflammatory Monocytes** - *Fcgr2b, H2-Eb1, Mrc1, Tap1, Tlr2* and *Tlr6*<br>**Macrophage** – *Ctss, Rab5c, Fcgr2b, H2-Q6, Sec22b, Dync1li1, Olr1, Itgav, Tubb5, Thbs1, Tuba1c, Tubb4b, Tlr2, C3, Cd14* and *Ctsl*<br>**Resident Macrophage** – *Marco, Olr1, Fcgr4, C3* and *Thbs1*<br>**Dendritic cell 1** – *Cd209a, Tap1, Olr1, Tlr2, Fcgr3, H2-DMb2, Fcgr4, Cybb* and *Fcgr1*<br>**Dendritic cell 2** – *H2-Q6, Tap1, H2-DMb2, Stx7, H2-Eb1, Atp6v0b, H2-Q7, H2-Aa, Cybb, H2-Ab1, Atp6v1g1, H2-DMa,* and *Coro1a*<br>**Dermal Dendritic cells** – *Msr1, Fcgr4, C3* and *Fcgr1* |
| Endocytosis | **Neutrophils** -*Cltc, Mdm2, Pdcd6ip, Rab11fip1, Chmp2a, Traf6, Smap1, Wipf1, Clta, Arfgef1, H2-K1, Smad3, Rab7, Rab11a, Arrb1, Rab31, Igf2r, Wipf2, Igf1r, Eps15, H2-T22, Hspa1b, Arpc4, Hspa8, Washc4, H2-T23, Ehd1, Kif5b, Vps4b, Hspa1a, Arf5, H2-Q10, Arf4, Arap1, Rab22a, Arpc2, Il2rg, H2-Q6, Arpc5, Psd4, Washc2, Tgfbr1, Stam2, H2-Q7, Cdc42, Capza2, Iqsec1, Eea1,* and *H2-D1*<br>**Inflammatory Monocytes** - *Dab2, Bin1, Igf1r, Ccr5, Arrb2,* and *Iqsec1*<br>**Macrophage** – *Itch, Rab5c, Cav2, Smurf1, Smurf2, Traf6, Wipf1, Src, Hspa1a, Snf8, Vps37b, Washc5, Snx12, Arfgef1, Chmp1a, Ldlrap1, Chmp1b, H2-Q6, Arrb1, Pip5k1a, Dab2, Capza1, Bin1, Dnm2, Igf1r, Snx1, Nedd4l, Stam2, Vps37a, Cav1, Gbf1, Agap1, Chmp5* and *Epn1*<br>**Resident Macrophage** – *Il2ra*<br>**Dendritic cell 1** – *Il2rg, Ccr5, Pip5k1c,* and *Cxcr4*<br>**Dendritic cell 2** – *Hspa1a, H2-Q6, Snx3, Rab8a, H2-Q7, Wipf1, Grk5, Capza2* and *Nedd4l*<br>**Dermal Dendritic cells** – *Cxcr2* |
| Efferocytosis | **Neutrophils** - *Hif1a, Sirpb1b, Rac1, Cd24a, Adam17, Itgav, Mapkapk2, Dusp16, Jak2, Sirpa, Tgfb1, Slc2a1, Ptpn6, Elmo1, Map2k1, Cebpb, Cd47, Tmem30a, Bsg, Pecam1, Rab7,* and *Lipa*<br>**Inflammatory Monocytes** - *Ptgs2, Hif1a, C1qb, Arg1, Cd24a,* and *Slc2a1*<br>**Macrophage** – *Ptgs2, Ptpn11, Mertk, Rab5c, Lrp1, Itgav, Thbs1, Mapk1, Mapk3, Arg2, Map2k1, Arg1, Sgk1, Arnt, Nfatc1, Bsg, Pecam1, Abca1, Ppard, Hif1a, Rab14, Cd24a, Slc2a1, Vps8, Atp11b, Sirpb1c,* and *Havcr2*<br>**Resident Macrophage** – *Arg2, Sirpb1b, Arg1, Sirpb1c, Sirpb1a, Thbs1, Slc2a1, Nr1h3, Sirpd,* and *Havcr2*<br>**Dendritic cell 1** – *Ptgs2, Hif1a, Arg1, Atp2a1, Adam17, Sirpb1c,* and *Slc2a1*<br>**Dendritic cell 2** – *Ptgs2, Hif1a, Arg2, Arg1,* and *Mapk3*<br>**Dermal Dendritic cells** – *Arg1, Sirpb1c,* and *Sirpd* |
| Ubiquitin mediated proteolysis | **Neutrophils** - *Wwp2, Herc4, Mdm2, Ube2a, Ube2g1, Fbxw11, Ube2n, Traf6, Cul3, Ube2d2a, Cul1, Ube2s, Ubc, Socs3, Ube2j1, Socs1, Ube2i, Birc6, Ube2z, Birc3, Herc1,* and *Xiap*<br>**Inflammatory Monocytes** - *Wwp2,* and *Socs1*<br>**Macrophage** – *Cul2, Itch, Wwp2, Herc4, Ube2g1, Ube2f, Smurf1, Smurf2, Ube2b, Traf6, Cul3, Cul4a, Ube3a, Ube2k, Cop1, Ube2l3, Cul5, Anapc16, Uba6, Map3k1, Uba1, Nedd4l, Pias1, Socs3,* and *Birc2*<br>**Dendritic cell 1** – *Ube2l6,* and *Anapc2*<br>**Dendritic cell 2** – *Nedd4l,* and *Socs3* |
| Fc gamma R-mediated phagocytosis | **Neutrophils** - *Arpc2, Prkcd, Pik3cb, Arpc5, Hck, Marcksl1, Rac1, Fcgr3, Inpp5d, Lyn, Vav1, Syk, Map2k1, Pik3r1, Vav3, Ncf1, Pik3ca, Cdc42, Arpc4,* and *Marcks*<br>**Inflammatory Monocytes** - *Fcgr2b, Pik3cb, Marcksl1, Bin1,* and *Pla2g4a*<br>**Macrophage** – *Fcgr2b, Pip5k1a, Pik3cb, Map2k1, Pik3r1, Bin1, Dnm2, Rac2, Pla2g4a, Was, Mapk1,* and *Mapk3*<br>**Resident Macrophage** – *Fcgr4*<br>**Dendritic cell 1** – *Ptprc, Fcgr3, Fcgr4, Rac2, Pip5k1c, Pik3cd,* and *Fcgr1* |

| | |
|---|---|
| | **Dendritic cell 2** – *Prkcd, Cfl1, Vasp, Mapk3,* and *Scin*<br>**Dermal Dendritic cells** – *Fcgr4,* and *Fcgr1* |
| Neutrophil extracellular trap formation | **Neutrophils** - *Fpr3, Rela, Pik3cb, Fpr1, C5ar1, Rac1, Casp4, H3f3a, Fcgr3, Syk, Tlr4, Map2k1, Pik3r1, Tlr2, C3, Ncf1, Fpr2, Pik3ca,* and *Cybb*<br>**Inflammatory Monocytes** - *Pik3cb, Fpr1, Tlr2,* and *Fpr2*<br>**Macrophage** – *Plcb1, Pik3cb, Hdac5, Hat1, Hdac1, Rac2, Mapk1, Src, Vdac3, Mapk3, Map2k1, Hdac4, Clcn3, Pik3r1, Tlr2, C3,* and *Fpr2*<br>**Resident Macrophage** – *Plcb2, Fpr1, Fcgr4, C3,* and *Itgal*<br>**Dendritic cell 1** – *Selplg, Gsdmd, Fcgr3, Tlr2, Fcgr4, Rac2, Fpr2, Cybb, Pik3cd,* and *Fcgr1*<br>**Dendritic cell 2** – *Casp4, Cybb, H3f3b,* and *Mapk3*<br>**Dermal Dendritic cells** – *Fpr3, Fpr1, Fcgr4, C3, Fpr2,* and *Fcgr1* |
| Toll-like receptor signaling pathway | **Neutrophils** - *Il1b, Casp8, Spp1, Jun, Nfkbia, Traf6, Tnf, Map2k1, Map3k8, Cd14, Cxcl10, Ccl3, Rela, Pik3cb, Rac1, Stat2, Stat1, Tlr6, Tlr4, Ccl4, Tbk1, Ikbke, Pik3r1, Tlr2, Irf5* and *Pik3ca*<br>**Inflammatory Monocytes** - *Ccl3, Il1b, Pik3cb, Spp1, Tlr2, Cd40, Ifnar1, Cxcl9, Stat2, Tlr6, Ccl5,* and *Ccl4*<br>**Macrophage** – *Jak1, Ccl3, Il1b, Pik3cb, Spp1, Traf6, Mapk1, Irf9, Mapk3, Map2k1, Pik3r1, Tlr2, Chuk, Ctsk,* and *Cd14*<br>**Resident Macrophage** – *Spp1* and *Cxcl9*<br>**Dendritic cell 1** – *Il1b, Irf7, Ccl3, Tlr2, Cd40, Pik3cd, Stat1* and *Ccl5*<br>**Dendritic cell 2** – *Il1b, Ccl3, Cd40, Cxcl9, Il12b, Mapk3* and *Ccl4*<br>**Dermal Dendritic cells** – *Tlr5, Spp1* and *Cxcl9* |
| C-type lectin receptor signaling pathway | **Neutrophils** - *Il1b, Casp8, Prkcd, Bcl3, Mdm2, Mapkapk2, Jun, Nfkbia, Tnf, Clec4e, Itpr2, Relb, Malt1, Irf1, Rela, Pik3cb, Clec4d, Stat2, Stat1, Syk, Ikbke, Pik3r1, Clec4n, Fcer1g, Pik3ca,* and *Nlrp3*<br>**Inflammatory Monocytes** - *Il1b, Ptgs2, Irf1, Pik3cb, Clec4e,* and *Stat2*<br>**Macrophage** – *Ptpn11, Il1b, Ptgs2, Malt1, Pik3cb, Kras, Clec4d, Mapk1, Src, Irf9, Mapk3, Pik3r1, Clec4n, Plk3, Chuk, Egr2, Nfatc1,* and *Itpr2*<br>**Dendritic cell 1** – *Cd209a, Il1b, Ptgs2, Malt1, Ccl17, Clec4n, Clec4e, Pik3cd, Stat1* and *Nlrp3*<br>**Dendritic cell 2** – *Il1b, Ptgs2, Prkcd, Clec4d, Ccl17, Il12b,* and *Mapk3*<br>**Dermal Dendritic cells** – *Calml4,* and *Ccl17* |
| NOD-like receptor signaling pathway | **Neutrophils** - *Il1b, Nfkbib, Casp8, Prkcd, Cxcl2, Il18, Aim2, Jun, Nfkbia, Rnasel, Traf6, Tnf, Cxcl3, Gbp7, Irgm2, Gbp5, Nampt, Irgm1, Nod1, Itpr2, Mcu, Ctsb, Rela, Ywhae, Casp4, Tank, Txn1, Stat2, Erbin, Stat1, Atg16l1, Tlr4, Tbk1, Ikbke, Hsp90aa1, Mefv, Gabarap, Tnfaip3, Ifi204, Bcl2l1, Cybb, Gbp3, Birc3, Gbp2, Nlrp3,* and *Xiap*<br>**Inflammatory Monocytes** - *Il1b, Cxcl2, Gbp5, Nampt, Ifnar1, Stat2, Gbp7, Cxcl3, Ccl5, Gbp2,* and *Irgm1*<br>**Macrophage** – *Jak1, Plcb1, Il1b, Map1lc3b, Cxcl2, Dnm1l, Traf6, Mapk1, Atg16l1, Irf9, Cxcl3, Vdac3, Mapk3, Gabarapl1, Cxcl1, Txnip, Atg5, Chuk, Birc2, Pkn1, Antxr2, Itpr2, Mcu,* and *P2rx7*<br>**Resident Macrophage** – *Plcb2* and *Trpm2*<br>**Dendritic cell 1** – *Il1b, Irf7, Gbp5, Gsdmd, Cybb, Stat1, Ccl5, Gbp2, Irgm1,* and *Nlrp3*<br>**Dendritic cell 2** – *Il1b, Prkcd, Gbp5, Cxcl2, Nampt, Gabarap, Casp4, Txn1, Sugt1, Cybb, Gbp2,* and *Mapk3* |
| HIF-1 signaling pathway | **Neutrophils** - *Hk3, Pgk1, Egln3, Map2k1, Pfkfb3, Eno1, Cdkn1a, Hmox1, Eif4e, Gapdh, Pfkl, Hif1a, Rela, Nos2, Pik3cb, Hk2, Igf1r, Hk1, Stat3, Slc2a1, Tlr4, Ldha, Pik3r1, Pfkp, Pik3ca, Aldoa,* and *Cybb*<br>**Inflammatory Monocytes** - *Hk3, Vegfa, Hif1a, Pik3cb, Nos2, Serpine1, Hk2, Igf1r, Hk1, Slc2a1,* and *Egln3*<br>**Macrophage** – *Cul2, Hk3, Egln1, Vegfa, Pgk1, Ltbr, Ifngr2, Mapk1, Egln3, Mapk3, Map2k1, Pfkfb3, Eno1, Cdkn1a, Eif4e, Gapdh, Arnt, Pfkl, Hif1a, Nos2, Pik3cb, Hk2, Igf1r, Hk1, Slc2a1, Ldha, Pik3r1, Trf, Aldoa,* and *Pdhb*<br>**Resident Macrophage** – *Nos2, Slc2a1* and *Egln3*<br>**Dendritic cell 1** – *Vegfa, Hif1a, Nos2, Pfkfb3, Cdkn1a, Cybb, Pik3cd, Slc2a1,* and *Egln3*<br>**Dendritic cell 2** – *Vegfa, Hif1a, Ldha, Hk2, Eno1, Gapdh, Cybb,* and *Mapk3*<br>**Dermal Dendritic cells** – *Nos2,* and *Egln3* |
| TNF signaling pathway | **Neutrophils** - *Il1b, Casp8, Bcl3, Cxcl2, Junb, Jun, Nfkbia, Tnf, Cxcl3, Map2k1, Csf1, Map3k8, Cxcl10, Cflar, Irf1, Rela, Pik3cb, Icam1, Tnfrsf1b, Atf2, Traf1, Atf4, Socs3, Pik3r1, Tnfaip3, Cebpb, Ifi47, Creb5, Pik3ca, Birc3,* and *Xiap* |

| | |
|---|---|
| | **Inflammatory Monocytes** - *Il1b, Ptgs2, Irf1, Pik3cb, Csf1, Cxcl2, Il15, Ifi47, Cxcl3,* and *Ccl5*<br>**Macrophage** – *Mmp14, Itch, Il1b, Ptgs2, Pik3cb, Cxcl2, Dnm1l, Atf2, Mapk1, Rps6ka5, Cxcl3, Mapk3, Cxcl1, Map2k1, Socs3, Pik3r1, Chuk, Birc2,* and *Jag1*<br>**Resident Macrophage** – *Mmp14* and *Vcam1*<br>**Dendritic cell 1** – *Il1b, Ccl5, Ptgs2, Ifi47,* and *Pik3cd*<br>**Dendritic cell 2** – *Il1b, Ptgs2, Socs3, Cxcl2, Creb5,* and *Mapk3* |
| NF-kappa B signaling pathway | **Neutrophils** - *Il1b, Cxcl2, Ltb, Nfkbia, Traf6, Tnf, Cxcl3, Ube2i, Cd14, Cflar, Relb, Malt1, Rela, Gadd45b, Icam1, Bcl2a1a, Gadd45a, Lyn, Tlr4, Traf1, Ccl4, Syk, Tnfaip3, Bcl2a1b, Bcl2l1, Bcl2a1d, Birc3,* and *Xiap*<br>**Inflammatory Monocytes** - *Il1b, Ptgs2, Cxcl2, Cd40, Cxcl3,* and *Ccl4*<br>**Macrophage** – *Il1b, Ptgs2, Malt1, Cxcl2, Ltbr, Gadd45a, Traf6, Cxcl3, Btk, Cxcl1, Chuk, Cd14,* and *Birc2*<br>**Resident Macrophage** – *Vcam1, Ltb,* and *Bcl2a1a*<br>**Dendritic cell 1** – *Il1b, Ptgs2, Malt1, Card11, Cd40,* and *Blnk*<br>**Dendritic cell 2** – *Il1b, Ptgs2, Cxcl2, Cd40* and *Ccl4* |
| Th1 and Th2 cell differentiation | **Neutrophils** - *Stat6, Nfkbib, Rbpj, Rela, Nfkbie, Il2rg, Jun, Jak2, Nfkbia, Il4ra, Stat1, H2-Aa,* and *H2-Ab1*<br>**Inflammatory Monocytes** – *Rbpj, H2-Eb1,* and *Il4ra*<br>**Macrophage** – *Jak1, Ifngr2, Il4ra, Chuk, Mapk1, Notch1, Nfatc1, Mapk3,* and *Jag1*<br>**Resident Macrophage** – *Il2ra, Il12rb2, Runx3, Cd3d,* and *Cd3e*<br>**Dendritic cell 1** – *Il2rg, Il12rb2, H2-DMb2,* and *Stat1*<br>**Dendritic cell 2** – *Il12rb2, H2-Eb1, H2-DMb2, Runx3, H2-Aa, H2-Ab1, Il12b, H2-DMa* and *Mapk3* |
| Th17 cell differentiation | **Neutrophils** – *Rara, Il1b, Nfkbib, Stat6, Rela, Hif1a, Nfkbie, Il2rg, Jun, Jak2, Runx1, Nfkbia, Il4ra, Stat3, Tgfb1, Stat1, Tgfbr1, Il1rap, Hsp90aa1, H2-Aa, H2-Ab1* and *Smad3*<br>**Inflammatory Monocytes** - *Il1b, Hif1a, H2-Eb1, Il4ra,* and *Il1rap*<br>**Macrophage** – *Il1b, Jak1, Il21r, Hif1a, Ifngr2, Il4ra, Chuk, Mapk1, Nfatc1, Mapk3,* and *Il1rap*<br>**Resident Macrophage** – *Il21r, Il2ra, Cd3d,* and *Cd3e*<br>**Dendritic cell 1** – *Il1b, Il21r, Hif1a, Il2rg, H2-DMb2,* and *Stat1*<br>**Dendritic cell 2** – *Il1b, Hif1a, H2-Eb1, H2-DMb2, H2-Aa, H2-Ab1, H2-DMa, Mapk3* and *Ahr*<br>**Dermal Dendritic cells** – *Il21r* |
| IL-17 signaling pathway | **Neutrophils** - *Il1b, S100a9, Rela, Casp8, Cxcl2, Jun, Nfkbia, Tnf, Traf6, Hsp90b1, Cxcl3, Tbk1, Ikbke, Lcn2, Hsp90aa1, Tnfaip3, Mapk6, Cebpb, Srsf1, Jund, Cxcl10,* and *Usp25*<br>**Inflammatory Monocytes** - *Il1b, Ptgs2, S100a8, Cxcl2,* and *Cxcl3*<br>**Macrophage** – *Il1b, Ptgs2, S100a9, Cxcl1, S100a8, Cxcl2, Srsf1, Chuk, Traf6, Mapk1, Cxcl3,* and *Mapk3*<br>**Resident Macrophage** – *Lcn2, S100a9,* and *S100a8*<br>**Dendritic cell 1** – *Il1b, Ptgs2, S100a9, S100a8,* and *Ccl17*<br>**Dendritic cell 2** – *Il1b, Ptgs2, S100a9, S100a8, Cxcl2, Ccl17,* and *Mapk3*<br>**Dermal Dendritic cells** – *Lcn2* and *Ccl17* |
| Chemokine signaling pathway | **Neutrophils** - *Nfkbib, Prkcd, Hck, Cxcl2, Pxn, Jak2, Nfkbia, Fgr, Cxcl3, Map2k1, Rap1a, Ncf1, Ptk2b, Cxcl10, Gnai3, Ccl3, Rela, Arrb1, Pik3cb, Rac1, Gnaq, Prex1, Stat3, Stat2, Lyn, Foxo3, Stat1, Ccl6, Vav1, Ccl4, Elmo1, Ccr1, Pik3r1, Vav3, Gsk3a, Gnb1, Pik3ca, Cdc42,* and *Rock2*<br>**Inflammatory Monocytes** - *Ccl3, Cxcl16, Pik3cb, Cxcl2, Stat2, Cxcl3, Ccl5, Ccl4, Pik3r6, Ccr9, Cxcl9, Ccr5,* and *Arrb2*<br>**Macrophage** – *Plcb1, Cxcl2, Pxn, Kras, Was, Mapk1, Src, Cxcl3, Mapk3, Map2k1, Chuk, Gngt2, Ccl3, Cxcl16, Arrb1, Pik3cb, Sos1, Rac2, Braf, Foxo3, Pik3r6, Ccr1, Cxcl1, Pik3r1, Prkacb,* and *Gng10*<br>**Resident Macrophage** – *Plcb2, Cxcl13, Cxcl9, Gngt2,* and *Fgr*<br>**Dendritic cell 1** – *Ccl3, Ccr1, Ccl17, Rac2, Ccr5, Pik3cd, Stat1, Fgr, Cxcr4, Ccl5,* and *Ccr7*<br>**Dendritic cell 2** – *Ccl3, Prkcd, Cxcl2, Gnb2, Ccl17, Foxo3, Ccl8, Fgr, Ccl4, Mapk3, Pik3r6, Cxcl9, Gngt2,* and *Grk5*<br>**Dermal Dendritic cells** – *Ccl17, Cxcl9, Gngt2,* and *Cxcr2* |
| Cytokine-cytokine receptor interaction | **Neutrophils** - *Il1b, Il1a, Cxcl2, Il18, Ltb, Il10rb, Il4ra, Tnf, Tgfb1, Cxcl3, Csf1, Cxcl10, Ccl3, Il13ra1, Il2rg, Inhba, Il15ra, Tnfrsf1b, Ccl6, Tgfbr1, Il1rn, Ccl4, Il1rap, Ccr1, Csf2rb,* and *Csf2rb2* |

| | |
|---|---|
| | **Inflammatory Monocytes** - *Ccl3, Il1b, Il7r, Il1a, Cxcl16, Cxcl2, Inhba, Il15, Il4ra, Cxcl3, Ccl5, Il1rn, Ccl4, Il1rap, Csf1, Ccr9, Cd40, Ifnar1, Cxcl9, Ccr5, Csf2rb,* and *Csf2rb2*<br>**Macrophage** – *Ccl3, Il1b, Il13ra1, Il7r, Cxcl16, Cxcl2, Inhba, Ltbr, Ifngr2, Il1r2, Il4ra, Cxcl3, Il1rn, Il1rap, Tnfrsf9, Il21r, Ccr1, Cxcl1, Tgfb3, Csf2rb,* and *Csf2rb2*<br>**Resident Macrophage** – *Il21r, Il18rap, Il12rb2, Il2ra, Ltb, Inhba, Cxcl13, Il1r2,* and *Cxcl9*<br>**Dendritic cell 1** – *Ccl3, Il1b, Il7r, Il2rg, Il12rb2, Osm, Inhba, Ccl17, Ccl5, Il1rn, Il21r, Ccr1, Cd40, Tnfsf4, Ccr5, Cxcr4,* and *Ccr7*<br>**Dendritic cell 2** – *Ccl3, Il1b, Il12rb2, Cxcl2, Acvr2a, Il1r2, Ccl17, Ccl8, Il12b, Il1rn, Ccl4, Cd40,* and *Cxcl9*<br>**Dermal Dendritic cells** – *Il21r, Osm, Inhba, Ccl17, Cxcl9, Csf3r,* and *Cxcr2* |
| JAK-STAT signaling pathway | **Neutrophils** - *Stat6, Jak2, Il10rb, Il4ra, Cdkn1a, Pim1, Il13ra1, Il2rg, Pik3cb, Il15ra, Ptpn2, Mcl1, Stat3, Stat2, Stat1, Ptpn6, Stam2, Socs3, Socs1, Pik3r1, Csf2rb, Bcl2l1, Pik3ca, Cish,* and *Csf2rb2*<br>**Inflammatory Monocytes** - *Il7r, Pik3cb, Socs1, Il15, Ifnar1, Il4ra, Csf2rb, Stat2,* and *Csf2rb2*<br>**Macrophage** – *Jak1, Ptpn11, Il13ra1, Il7r, Pik3cb, Sos1, Ifngr2, Ptpn2, Il4ra, Irf9, Stam2, Il21r, Pias1, Socs3, Pik3r1, Cdkn1a, Csf2rb, Socs4, Cish,* and *Csf2rb2*<br>**Resident Macrophage** – *Il21r, Il2ra,* and *Il12rb2*<br>**Dendritic cell 1** – *Il21r, Il7r, Il2rg, Il12rb2, Osm, Cdkn1a, Pik3cd, Stat1,* and *Cish*<br>**Dendritic cell 2** – *Socs3, Il12rb2,* and *Il12b*<br>**Dermal Dendritic cells** – *Il21r, Osm,* and *Csf3r* |
| Arginine biosynthesis | **Neutrophils** - *Nos2*<br>**Inflammatory Monocytes** - *Got1, Nos2,* and *Arg1*<br>**Macrophage** – *Got1, Nos2, Arg2, Arg1,* and *Ass1*<br>**Resident Macrophage** – *Nos2, Arg2, Arg1*, and *Ass1*<br>**Dendritic cell 1** – *Nos2* and *Arg1*<br>**Dendritic cell 2** – Got1, Arg2, Arg1, and Ass1<br>**Dermal Dendritic cells** – *Nos2* and *Arg1* |
| Antigen processing and presentation | **Neutrophils** - *Ctss, Hspa8, H2-T23, Psme2, Canx, B2m, Tnf, Psme1, Hspa1a, H2-Q10, H2-K1, H2-Q6, Ctsb, Tapbp, Tap2, Tap1, Hspa5, Hsp90aa1, H2-Q7, H2-T22, Cd74, Hspa1b, H2-Aa, H2-Ab1,* and *H2-D1*<br>**Inflammatory Monocytes** - *Tap1,* and *H2-Eb1*<br>**Macrophage** – *Ctss, Hspa1a, H2-Q6, Ifi30, Nfya,* and *Ctsl*<br>**Dendritic cell 1** – *Cd8b1, Tap1, Ifi30, H2-DMb2,* and *Cd8a*<br>**Dendritic cell 2** – *Hspa1a, H2-Q6, H2-Eb1, Tap1, Psme2, H2-DMb2, H2-Q7, Ciita, H2-Aa, Cd74, H2-Ab1*, and *H2-DMa* |
| Complement and coagulation cascades | **Neutrophils** - *C5ar1, Plaur, F3, C3,* and *F10*<br>**Inflammatory Monocytes** - *C1qb, Serpine1, F3, Cfb, F13a1, F10,* and *C3ar1*<br>**Macrophage** – *Procr, Plaur, Cfb, C3, F10, Thbd,* and *Serpinb2*<br>**Resident Macrophage** – *F7, C3* and *F10*<br>**Dendritic cell 1** – *F10*<br>**Dendritic cell 2** – *Procr*<br>**Dermal Dendritic cells** – *F7, C3* and *Cfb* |
| PI3K-Akt signaling pathway | **Neutrophils** - *Mdm2, Spp1, Itgav, Jak2, Il4ra, Ywhaz, Map2k1, Csf1, Ppp2r2d, Cdkn1a, Eif4e, Itgb1, Pik3ap1, Lamb3, Rela, Il2rg, Pik3cb, Ddit4, Ywhae, Rac1, Igf1r, Pten, Mcl1, Ppp2ca, Crtc2, Atf2, Foxo3, Hsp90b1, Tlr4, Syk, Atf4, Hsp90aa1, Pik3r1, Ppp2r1a, Tlr2, Gnb1, Creb5, Bcl2l1,* and *Pik3ca*<br>**Inflammatory Monocytes** - *Vegfa, Il7r, Pik3cb, Spp1, Met, Igf1r, Il4ra, Pik3r6, Csf1, Gys1, Tlr2, Ifnar1,* and *Lpar1*<br>**Macrophage** – *Jak1, Vegfa, Il7r, Pgf, Spp1, Kras, Itgav, Il4ra, Thbs1, Lamc1, Mapk1, Ppp2r5e, Mapk3, Map2k1, Gys1, Cdkn1a, Sgk1, Chuk, Gngt2, Eif4e, Pkn1, Ppp2r2a, Itga1, Pik3cb, Ddit4, Sos1, Met, Itgb7, Igf1r, Pten, Atf2, Foxo3, Pik3r6, Pik3r1, Tlr2, Bcl2l11, Pdpk1, Gng10,* and *Cdk4*<br>**Resident Macrophage** – *Pgf, Il2ra, Spp1, Hgf, Gngt2,* and *Thbs1*<br>**Dendritic cell 1** – *Vegfa, Il7r, Il2rg, Ddit4, Pgf, Gys1, Osm, Tlr2, Areg, Cdkn1a,* and *Pik3cd*<br>**Dendritic cell 2** – *Vegfa, Pik3r6, Ddit4, Ywhag, Gnb2, Creb5, Gngt2, Bcl2l11, Foxo3, Ywhaq, Kit,* and *Mapk3*<br>**Dermal Dendritic cells** – *Gys1, Osm, Spp1, Gngt2,* and *Csf3r* |

**Table S4:** The DEGs of lymphoid subsets enriched in the KEGG pathways upon *C. auris* murine skin infection.

| KEGG Pathways | DEGs enriched in the pathway |
|---|---|
| Th17 cell differentiation | **CD4+ Th Cells** - *Il23r, Stat5b, Hif1a, Stat5a, Il2ra, Il22, Il1r1, Nfkbia, Il17a, Lck, Cd4, Ifng, Cd247,* and *Cd3d*<br>**CD8+ Cells** – *Nfkbia*<br>**γδ+ T cells** – *Hif1a, Il22,* and *Ifng* |
| Th1 and Th2 cell differentiation | **CD4+ Th Cells** – *Lck, Stat5b, Stat5a, Cd4, Il2ra, Nfkbia, Cd247, Ifng* and *Cd3d*<br>**CD8+ Cells** – *Nfkbia*<br>**γδ+ T cells** – *Il13, Il12rb2,* and *Ifng* |
| IL-17 signaling pathway | **CD4+ Th Cells** – *Nfkbia, Csf2, Ifng, Il17a,* and *Cxcl3*<br>**CD8+ Cells** – *S100a9, S100a8, Cxcl2* and *Nfkbia*<br>**γδ+ T cells** – *Ptgs2, Il13, Csf2,* and *Ifng* |
| TNF signaling pathway | **CD4+ Th Cells** - *Csf1, Nfkbia, Csf2, Cxcl3* and *Cflar*<br>**Tregs** - *Map2k1*<br>**CD8+ Cells** – *Cxcl2* and *Nfkbia*<br>**γδ+ T cells** – *Ptgs2, Csf2, Ccl5* and *Cflar* |
| T cell receptor signaling pathway | **CD4+ Th Cells** – *Lck, Cd4, Nfkbia, Ctla4, Csf2, Nck2, Cd247, Ifng,* and *Cd3d*<br>**Tregs** - *Map2k1*<br>**CD8+ Cells** – *Icos,* and *Nfkbia*<br>**γδ+ T cells** – *Ctla4, Csf2,* and *Ifng* |
| B cell receptor signaling pathway | **CD4+ Th Cells** - *Lilrb4a, Ifitm1, Nfkbia,* and *Pik3ap1*<br>**Tregs** - *Map2k1* and *Prkcb*<br>**CD8+ Cells** – *Nfkbia* |
| Chemokine signaling pathway | **CD4+ Th Cells** - *Stat5b, Ccr1, Plcb4, Nfkbia, Gnaq,* and *Cxcl3*<br>**Tregs** – *Prkcb, Map2k1,* and *Ccl4*<br>**CD8+ Cells** – *Cxcl2, Nfkbia,* and *Ccl4*<br>**γδ+ T cells** – *Ccr5* and *Ccl5* |
| Cytokine-cytokine receptor interaction | **CD4+ Th Cells** - *Il23r, Stat5b, Hif1a, Stat5a, Il2ra, Il22, Il1r1, Nfkbia, Il17a, Lck, Cd4, Ifng, Cd247,* and *Cd3d*<br>**Tregs** - *Il1r2* and *Ccl4*<br>**CD8+ Cells** –*Cxcl2, Il1rn,* and *Ccl4*<br>**γδ+ T cells** – *Il13, Il18rap, Il12rb2, Il22, Il1r2, Ccr5, Csf2, Ifng,* and *Ccl5* |
| JAK-STAT signaling pathway | **CD4+ Th Cells** - *Il23r, Stat5b, Stat5a, Il2ra, Il22, Csf2, Pim1,* and *Ifng*<br>**γδ+ T cells** – *Il13, Il12rb2, Il22, Pdgfb, Csf2,* and *Ifng* |
| NF-kappa B signaling pathway | **CD4+ Th Cells** – *Lck, Ltb, Il1r1, Nfkbia, Cxcl3,* and *Cflar*<br>**Tregs** - *Prkcb* and *Ccl4*<br>**CD8+ Cells** – *Cxcl2, Ccl4* and *Nfkbia*<br>**γδ+ T cells** – *Cflar* and *Ptgs2* |
| HIF-1 signaling pathway | **CD4+ Th Cells** - *Pfkl, Egln1, Hif1a, Ldha, Pgk1, Pfkp, Eno1,* and *Ifng*<br>**Tregs** - *Egln1, Prkcb, Map2k1, Hk2* and *Prkca*<br>**γδ+ T cells** – *Hif1a, Eno1,* and *Ifng* |
| VEGF signaling pathway | **CD4+ Th Cells** - *Mapkapk2*<br>**Tregs** – *Prkcb, Map2k1* and *Prkca*<br>**γδ+ T cells** – *Ptgs2* and *Sh2d2a* |
| Wnt signaling pathway | **CD4+ Th Cells** - *Plcb4, Lef1,* and *Ccn4*<br>**Tregs** – *Prkcb, Prkca, Tbl1x,* and *Lef1* |
| Rap1 signaling pathway | **CD4+ Th Cells** - *Csf1, Plcb4, Gnaq, Rapgef1,* and *Itgal*<br>**Tregs** – *Prkcb, Map2k1,* and *Prkca*<br>**CD8+ Cells** – *Thbs1* |

| | |
|---|---|
| | **γδ+ T cells** – *Pdgfb*<br>**NK Cells** – *Evl* |
| Fc epsilon RI signaling pathway | **CD4+ Th Cells** – *Csf2*<br>**Tregs** - *Map2k1*, and *Prkca*<br>**γδ+ T cells** – *Csf2* and *Il13* |
| Fc gamma R-mediated phagocytosis | **CD4+ Th Cells** - *Myo10*, and *Actr3*<br>**Tregs** – *Prkcb*, *Map2k1*, and *Prkca*<br>**γδ+ T cells** – *Myo10* |
| Natural killer cell mediated cytotoxicity | **CD4+ Th Cells** – *Lck*, *Klrc1*, *Csf2*, *Itgal*, *Cd247*, and *Ifng*<br>**Tregs** – *Prkcb*, *Map2k1*, *Gzmb*, and *Prkca*<br>**γδ+ T cells** – *Klrc2*, *Klrc1*, *Csf2*, and *Ifng* |

**Table S5:** The DEGs of fibroblast subsets enriched in the KEGG pathways upon *C. auris* murine skin infection.

| KEGG Pathways | DEGs enriched in the pathway |
|---|---|
| HIF-1 signaling pathway | **Fibroblast 1** - *Nos2, Serpine1* and *Timp1*<br>**Fibroblast 3** – *Igf1, Pfkl, Nos2, Serpine1, Elob, Eno1, Eif4ebp1, Timp1, Slc2a1* and *Angpt1* |
| PI3K-Akt signaling pathway | **Fibroblast 1** - *Ddit4, Pgf, Ereg, Col6a5, Tnc, Pik3ap1,* and *Csf3*<br>**Fibroblast 2**- *Csf3r*<br>**Fibroblast 3** – *Igf1, Ddit4, Pgf, Spp1, Itga11, Fgf23, Thbs4, Thbs3, Tnc, Creb3l3, Col6a2, Gys1, Ereg, Eif4ebp1, Tnn, Angpt1,* and *Csf3* |
| ECM-receptor interaction | **Fibroblast 1** - *Col6a5* and *Tnc*<br>**Fibroblast 3** – *Col6a2, Spp1, Itga11, Thbs4, Thbs3, Tnn,* and *Tnc* |
| IL-17 signaling pathway | **Fibroblast 1** - *Lcn2, S100a9, S100a8, Mmp13, Cxcl3,* and *Csf3*<br>**Fibroblast 2**- *S100a9, S100a8, Cxcl2,* and *Cxcl3*<br>**Fibroblast 3** – *Lcn2, S100a9, Cxcl5, Mmp13,* and *Csf3*<br>**Fibroblast 4** – *S100a9* |
| NF-kappa B signaling pathway | **Fibroblast 1** - *Cxcl3*<br>**Fibroblast 2**- *Cxcl3, Btk* and *Cxcl2*<br>**Fibroblast 3** – *Lbp* and *Cxcl12* |
| Cytokine-cytokine receptor interaction | **Fibroblast 1** - *Il13ra2, Il33, Cxcl9, Tnfsf8, Cxcl3,* and *Csf3*<br>**Fibroblast 2**- *Cxcl2, Csf3r,* and *Cxcl3*<br>**Fibroblast 3** – *Cxcl5, Cxcl14, Il33, Il11ra1, Bmpr1b, Tnfsf8, Il18r1, Cxcl12* and *Csf3* |
| TNF signaling pathway | **Fibroblast 1** - *Cxcl3*<br>**Fibroblast 2**- *Cxcl3,* and *Cxcl2*<br>**Fibroblast 3** – *Cxcl5, Creb3l3* and *Il18r1* |
| Chemokine signaling pathway | **Fibroblast 1** - *Cxcl9* and *Cxcl3*<br>**Fibroblast 2**- *Cxcl3* and *Cxcl2*<br>**Fibroblast 3** – *Cxcl5, Cxcl14* and *Cxcl12* |
| NOD-like receptor signaling pathway | **Fibroblast 1** - *Cxcl3*<br>**Fibroblast 2**- *Cxcl3* and *Cxcl2*<br>**Fibroblast 3** – *Irf7* |
| Complement and coagulation cascades | **Fibroblast 1** - *Kng2, Serpine1, Cfb, Cfi,* and *Serpinb2*<br>**Fibroblast 3** – *Bdkrb2, Serpine1, Cfb, C3,* and *Serpinb2* |

**Table S6:** The list of antibodies, ELISA kit, and recombinant proteins used in the study.

| Reagent or Resource | Source | Identifier |
|---|---|---|
| **Antibodies** | | |
| Anti-mouse CD11c, APC (N418) | Biolegend | Cat# 117309; RRID: AB_313779 |
| Anti-mouse Ly6C, Pacific Blue (HK1.4) | Biolegend | Cat# 128014; RRID: AB_1732079 |
| Anti-mouse CD11b, PE (M1/70) | Biolegend | Cat# 101208; RRID: AB_312791 |
| Anti-mouse Ly6G, PE/Cyanine7 (1A8) | Biolegend | Cat# 127618; RRID: AB_1877261 |
| Anti-mouse Ly6G, APC (S19018G) | Biolegend | Cat#164506; RRID: AB_2927993 |
| Anti-mouse CD45, FITC (30-F11) | Biolegend | Cat# 103108; RRID: AB_312973 |
| Anti-mouse CD64, PE/Dazzle 594 (X54-5/7.1) | Biolegend | Cat# 139320; RRID: AB_2566559 |
| Anti-mouse MHC II, Alexa Fluor 700 (M5/114.15.2) | eBioscience | Cat# 56532182; RRID: AB_494009 |
| Anti-mouse TCR γ/δ, BV421 (GL3) | Biolegend | Cat# 118120; RRID: AB_2562566 |
| Anti-mouse CD4, PerCP/Cyanine5.5 (GK1.5) | Biolegend | Cat# 100434; RRID: AB_893324 |
| Anti-mouse CD8b, PE (H35-17.2) | BD Biosciences | Cat# 550798; RRID: AB_393887 |
| Anti-mouse TCR β, PE/Cyanine7 (H57-597) | Biolegend | Cat# 109222; RRID: AB_893625 |
| Anti-mouse IL-17F, Alexa Fluor 488 (9D3.1C8) | Biolegend | Cat# 517006; RRID: AB_10661903 |
| Anti-mouse IL-17A, PE/Dazzle 594 (TC11-18H10.1) | Biolegend | Cat# 506938; RRID: AB_2564321 |
| Anti-mouse IFN-γ, Alexa Fluor 700 (XMG 1.2) | Biolegend | Cat# 505824; RRID: AB_2561300 |
| Anti-mouse CD16/32 (93) | Biolegend | Cat# 101302; RRID: AB_312801 |
| Anti-mouse F4/80, PE/Dazzle 594 (BM8) | Biolegend | Cat# 123145; RRID: AB_2564132 |
| Anti-mouse F4/80, Alexa Fluor 700 (BM8) | Biolegend | Cat# 123129; RRID: AB_2277848 |
| Ultra-LEAF™ Purified anti-mouse Ly-6G Antibody (1A8) | Biolegend | Cat# 127649; RRID: AB_2572001 |
| Ultra-LEAF™ Purified Rat IgG2a, κ Isotype Ctrl Antibody (RTK2758) | Biolegend | Cat# 400565; RRID: AB_11147167 |
| Anti-mouse IL-1Ra monoclonal antibody | This study | Gift from Naofumi Mukaida |
| **ELISA kit** | | |
| Mouse IL-1ra/IL-1F3 DuoSet ELISA | RnD | Cat# DY480 |
| **Recombinant Protein** | | |
| Recombinant Mouse IL-1RA (IL-1RN) | Biolegend | Cat# 769704 |

**Table S7:** The oligonucleotide sequences used to construct *pmr1* deletion and the confirmation of the desired edit by Sanger sequencing.

| **Description** | Sequence (5' → 3') |
|---|---|
| B9J08_000837 (*pmr1*) deletion gRNA | TATCGGAAAAGAACCCGTCG |
| Fragment 1 Forward Primer | TGGCGCTCTAGCACATTACC |
| Fragment 1 Reverse Primer | CCTCATGTCGAGCACTCGTCTCGCCGAGGATATGTATTAGGGG |
| Fragment 2 Forward Primer | CGAGACGAGTGCTCGACATGAGGTAGCCCACTTCATGTTGAA |
| Fragment 2 Reverse Primer | CCTCCAGTTTCTACAGAACAGGTGG |
| Colony PCR Forward Primer | CGCTCCGCCATATAACCCAT |
| Colony PCR Reverse Primer | ACGTAACTAGCCTTGACGGG |
| Sanger sequencing confirmation of *pmr1Δ* stain | NNNNNNNNTCTNGTTTGNGGTGCTCGCGCTCCGCCATATAAC CCATGAATCCTTCATCGATTCCACCTGGGGTCCCTATA TCTCCCAAACCACTAGCATCCCTCGCCCACAACCCAACCCCC TAATACATATCCTCGGCGAGACGAGTGCTCGACATGAG GTAGCCCACTTCATGTTGAACAGCCCCGTCAAGGCTAGTTACGT ATATATTATTTAGAGATTTCTGTCTAGAAATCGACA ATATCATCAAGACTAACAACGCCATTGGCTTCTGGGTCCTCAAG CTTGCGATAAAGCGTGATGGCCACCTGTTCTGNNNA ANCTGGNNGN |

**Table S8:** The fungal strains, plasmids and mouse strains used in the study.

| **Fungal Strains** | | |
|---|---|---|
| **Strains** | **Source** | **Genotype** |
| *Candida auris* AR0387 | CDC | Wild type |
| *Candida auris* AR0381 | CDC | Wild type |
| *Candida auris* AR0383 | CDC | Wild type |
| *Candida auris* AR0385 | CDC | Wild type |
| *C. auris* AR0387 *pmr1Δ* | This Study | *pmr1Δ* |
| *Candida albicans* SC5314 | Gifted from Dr. Andrew Koh | Wild type |
| **Experimental Models: Organisms/Strains** | | |
| **Strains** | **Source** | **Reference identifier** |
| C57BL/6J mice | The Jackson Laboratory | Cat#000664; RRID: IMSR_JAX:000664 |
| *IL-1R1*$^{-/-}$ C57BL/6J mice | The Jackson Laboratory | Cat# 003245 RRID: IMSR_JAX:003245 |
| **Plasmids** | **Description** | **Identifier** |
| pCE35 | CAS9 expression cassette | #174409 |
| pCE27 | gRNA expression cassette | #174405 |

**Table S9:** The list of reagents and software used in this study

| **Chemicals, Peptides, and Recombinant Proteins** | | |
|---|---|---|
| Yeast Extract-Peptone-Dextrose (YPD) | BD Bioscience | Cat # 242810 |
| Recombinant Mouse IL-1RA (IL-1RN) | Biolegend | Cat# 769704 |
| Agar | Fisher Scientific | Cat # BP1423-500 |
| 10× Phosphate Buffer Solution (PBS) | Fisher Scientific | Cat # BP3994 |
| Ampicillin | Cayman Chemicals | Cat # 14417 |
| Streptomycin | MP Biomedicals | Cat # 100556 |
| Bovine Serum Albumin (BSA) | Sigma Aldrich | Cat # A4737 |
| RPMI 1640 Medium (1X) with Glutamine and Phenol Red | ThermoFisher | Cat # 11875093 |
| Fetal Bovine Serum (FBS) | CPS Serum | Cat #: FBS-500HI |
| Percoll | Sigma Aldrich | Cat # P1644 |
| Triton X-100 | MP Biomedicals | Cat # 807423 |
| HyClone RPMI 1640 media with L-glutamine | Cytiva | Cat # SH30027.02 |
| Trypsin-EDTA (0.25%), phenol red | ThermoFisher | Cat # 25200056 |
| Cell Staining Buffer | Biolegend | Cat # 420201 |
| Intracellular Staining Permeabilization Wash Buffer | Biolegend | Cat # 421002 |
| Fixation Buffer | Biolegend | Cat # 420801 |
| Monensin Solution (1,000X) | Biolegend | Cat # 420701 |
| Cell Activation Cocktail (without Brefeldin A) | Biolegend | Cat # 423302 |
| LIVE/DEAD™ Fixable Yellow Dead Cell Stain kit, (405 nm) excitation | Invitrogen, ThermoFisher | Cat # L34959 |
| Liberase TL Research Grade | Sigma Aldrich | Cat # 05401020001 |
| Heparinized capillary tubes | Fisher Scientific | Cat # 22-260950 |
| Deoxyribonuclease I from bovine pancreas | Sigma Aldrich | Cat # DN25-100G |
| Sterile Cell Strainer, 70 µm | Fisherbrand | Cat# 22-363-548 |

| | | |
|---|---|---|
| Sterile Cell Strainer, 40 µm | Fisherbrand | Cat# 22-363-547 |
| MACS Smart Strainers | Miltenyi Biotec | Cat# 130-098-458 |
| **Software and Algorithms** | | |
| FlowJo v9 | Tree Star | https://www.flowjo.com/solutions/flowjo/downloads |
| GraphPad Prism 4 | GraphPad Software | https://www.graphpad.com/scientificsoftware/prism/ |
